# Microsporidia Ser/Thr Protein Phosphatase PP1 Targets DCs MAPK Pathway and Impairs Immune Functions

**DOI:** 10.1101/2023.09.13.557560

**Authors:** Jialing Bao, Yunlin Tang, Yebo Chen, Jiangyan Jin, Xue Wang, Guozhen An, Lu Cao, Biying Mo, Huarui Zhang, Gong Cheng, Guoqing Pan, Zeyang Zhou

**Affiliations:** The State Key Laboratory of Resource Insects, Southwest University, Chongqing, 400715, China; Chongqing Key Laboratory of Microsporidia Infection and Control, Southwest University, Chongqing 400715, China; Tsinghua University-Peking University Joint Center for Life Sciences, School of Medicine, Tsinghua University, Beijing 100084, China

**Keywords:** Microsporidia, *Enterocytozoon hellem*, Dendritic cells, Serine/Threonine Protein Phosphatase, MAPK

## Abstract

Microsporidia are difficult to completely eliminate. Their persistence may disrupt host cell functions. Here in this study, we aimed to elucidate the impairing effects and consequences of microsporidia infection upon dendritic cells (DCs). We used the zoonotic microsporidia species, *Enterocytozoon hellem*, in our studies. *In vivo* experiments showed that *E. hellem-*infected mice were more susceptible to further pathogenic challenges. DCs were identified as the most affected group of cells. *In vitro* assays revealed that *E. hellem* infection impaired the immune functions of DCs as reflected by down-regulation of cytokine expression, lower extent of maturation and antigen presentation. *E. hellem* infection decreased the ability of DCs to prime and stimulate T cells, thereby hampering host immune cell functions. We further demonstrate that *E. hellem* Ser/Thr protein phosphatase PP1 directly interacts with host p38α(MAPK14) to manipulate the p38α (MAPK14)/NFAT-5 axis of the MAPK pathway. Our study is the first to elucidate the molecular mechanisms of the impairing effects of microsporidia on host DCs immune functions. The emerging of microsporidiosis may be great threat to public health.

**Highlights:**

- Persistence of Microsporidia within host impairs dendritic cell functions such as phagocytosis, maturation, antigen presentation and T cell priming, thereby disrupting both innate and adaptive immunities and making the host more vulnerable to secondary infections
- Microsporidia impairs DCs function via Serine/Threonine Protein Phosphatase PP1 directly targeting DCs p38α/MAPK pathway
- Latent Microsporidia infection and persistence is a great threat to public health when assessing acute and emerging pathogen risk

## Introduction

Microsporidia ares a large group of intracellular pathogens that can infect all animals, from invertebrates to vertebrates including human beings (1, 2). At least 15 species can infect humans, with the four most common ones being: *Enterocytozoon bieneusi* (*E. bieneusi*), *Encephalitozoon hellem* (*E. hellem*), *Encephalitozoon cuniculi* (*E. cuniculi*) and *Encephalitozoon intestinalis* (*E. intestinalis*) (3, 4). Immune-compromised individuals are thought to be more vulnerable to microsporidia infections (5). However, accumulating evidence showsthat immune-competent individuals can also be infected, awith infection outcomes often asymptomatic and latent (6–8). For instance, one study conducted in Cameroon reveals that 87% teenagers and 68% healthy asymptomatic individuals tested actually had subclinical microsporidial infections and were shedding spores (9). In addition, co-infections of microsporidia with other pathogens such as HIV, cryptosporidia and *M. tuberculosis* are under-estimated in humans and patents usually exhibited exacerbated outcomes compared to single pathogen infection alone (10, 11). These findings reveal that latent infection and persistence of microsporidia in immune-competent individuals are much higher than previously thought. Although the levels of latent infection of microsporidia in immune-competent individuals have long been under-estimated, it should now be paid more attention to due to wide existence of microsporidia in nature and their wide host range them may lead to emerging infections and serious public health problems (8, 12).

Since microsporidia is an obligate intracellular pathogen,its interactions with host cells should be spotlighted, we are very interested to elucidate how microsporidia modulate host cell functions, especially the immune cell functions. Thus far, most studies were carried out using genetically knock-out or immune-deficient mice to increase microsporidia infection and colonization rates, and usually focused on host adaptive immunity (13, 14). There is an unmet demand to use these sameimmune-competent animal models to elucidate the roles of host sentinel innate immunity during microsporidia-host interactions (15, 16).

Dendritic cells (DCs), the professional antigen presenting cells and the bridge of innate and adaptive immunities, are found to participate in defending against microsporidia infections (17–19). Moretto *et al* showed that *in vitro* infection of *E. cuniculi* to DCs can stimulate CD8+ T cells to release IFNγ which is known to be cytotoxic to microsporidia; Bernal *et al* found that *E. intestinalis* interferes with DC differentiation and DC production of pro-inflammatory cytokine IL-6 (14, 15). Moreover, studies revealed that DCs differentiation influences host’s overall anti-microbial capability (20, 21). Therefore, DCs might be the master regulator of host immune responses against microsporidia and other pathogen co-infections.

The MAPK signaling pathway is essential in regulating many cellular processes including inflammation and stress responses. Along the signal transduction route, there are several key proteinases that can be modulated thus affect signaling outcomes. For example, the p38a (MAPK14), MAP Kinase Kinases (MKKs) have been reported to be regulated by several pathogens and medicines (22, 23). In addition, downstream of MAPK pathway, there is an essential transcriptional factor NFAT5 (24, 25). NFAT (Nuclear Factors of activated T cells) was originally identified as a key transcription factor involved in maintaining cellular homeostasis against hypertonic. Emerging evidence pointed out the immune-regulatory function of NFAT5, and is achieved by inducing different target genes and different signaling pathways. Expressions of pro-inflammatory genes such as *IL-6, IL-2*, *H2Ab* are all directly controlled by NFAT5 (26, 27).

Ubiquitous serine/threonine protein phosphatase (PP1) is a single domain catalytic protein that is exceptionally well conserved in all eukaryotes, from fungi to human, in both sequence and function (28). In humans, PP1 is responsible for about 30% of all de-phosphorylation reactions (29, 30). Pathogenic microbes usually express the serine/threonine proteinase as modulator of host. For example, *Mycobacterium tuberculosis* express Ser/Thr proteinase PknG within host macrophages to regulate host protein phosphorylation and to interfere with autophagy flux, thus greatly affect host cell functions (31, 32). Pathogenic microsporidia express only a subset of their genome, the bare minimum of their essential genes, during intracellular infection. However, we are very excited to have found the existence of *PP1* gene in microsporidia genome, indicating its key functions in pathogen growth and in pathogen-host interactions. It is therefore of great interest to exploit the regulation effects of microsporidia PP1 on dendritic cell functions.

Here in our study, we planned to utilize murine models, as well as cells cultured *in vitro,* to thoroughly investigate the effects of microsporidia infection on DCs. Our study will elucidate the regulation mechanisms of microsporidia on host immune responses, and will shed light on prevention of pathogens co-infections and emerging diseases.

## Materials and Methods

### Pathogens

*Encephalitozoon hellem* (*E. hellem*) strain (ATCC 50504) were a gift from Prof. Louis Weiss (Albert Einstein College of Medicine, USA). Spores were inoculated and propagated in rabbit kidney cells (RK13, ATCC CCL-37), cultured in Minimum Essential Medium Eagle (MEM) with 10% fetal bovine serum (FBS) (Thermo Fisher, USA). The spores were collected from culture media, purified by passing them through 5 μm size filter (Millipore, Billerica, MA), and stored in sterile distilled water at 4°C (33). Spores were counted with a hemocytometer before usages.

*Staphylococcus aureus* (S. aureus) was gifted by Dr. Xiancai Rao (Department of Microbiology, College of Basic Medical Sciences, Army Medical University, Chongqing, China). The microbe was modified on the basis of strain of N315 to express EGFP for visibility during microscopy, and cultured on TSB medium (34).

### Animals

Wild-type C57BL/6 mice (six-week, female) were reared in animal care facility according to Southwest University-approved animal protocol (SYXK-2017-0019). At the end of experiment, all mice were euthanized using carbon dioxide narcosis and secondary cervical dislocation.

### Primary cells and cell lines

Primary dendritic cells were isolated from mouse mesenteric lymph nodes, spleen and bone marrow. Mesenteric lymph nodes associated with the intestine were moved by forceps, washed with cold PBS and teased into single cell mix in harvesting medium such as RPMI Medium 1640. Bone marrow was flushed by RPMI Medium 1640 from the femurs of dissected mice. Spleens were digested in spleen dissociation medium (STEMCELL Technologies, Canada) at room temperature, followed by gently passing several times through a 16 Gauge blunt-end needle attached to a 3cc syringe and then passed through a primed 70μm nylon mesh filter. The single cell suspension, either from bone marrow or spleen, was then subjected to the EasySep Mouse Pan-DC Enrichment Kit (STEMCELL Technologies, Canada) to isolate dendritic cells. Briefly Enrichment Cocktail and subsequently Biotin Selection Cocktail were added to samples. Next, samples were incubated with magnetic particles and dendritic cells were selected via negative selection using magnetic separation. The isolated dendritic cells were then counted and cultured in RPMI Medium 1640 (supplemented with 10% FBS and penicillin/streptomycin) (Gibco, USA) in a 37°C, 5% CO2 incubator.

The dendritic cell line, DC2.4 (Sigma-Aldrich SCC142), was purchased from BeNa Culture Collection, China. The cells were cultured in RPMI Medium 1640 supplemented with 10% FBS and penicillin/streptomycin (Gibco, USA) in a 37°C, 5% CO_2_ incubator.

The suspension T-cell line, Jurkat cells (Clone E6-1, ATCC) was purchased from FuDan IBS Cell Center, China. The cells were cultured in RPMI Medium 1640, supplemented with 10% (v/v) FBS and 1% (v/v) penicillin/streptomycin (Gibco, USA) in a 37°C, 5% CO2 incubator.

### Microsporidia infection

*In vivo* infection of microsporidia of wild-type mice was achieved as follows. 1 × 10^7^ *E. hellem* spores/mice/day were inoculated into wild-type mice for two days; the mice were also transiently pre-treated with dexamethasone (Aladin, Cas 2392-39-4, China) to increase infection rates, we noted no significant effects on immune cells as assessed in our established murine model and other reports (35–38). The control groups were either treated with PBS, LPS (5mg/kg), or *S. aureus* (1× 10^7^ CFU/mice). Mouse body weights were monitored throughout infection time courses and at experimental endpoints, mice were sacrificed by CO_2_ inhalation and secondary cervical dislocation. Samples of mouse blood, urine, faeces and organs were also collected for further experiments.

*In vitro* infection of microsporidia was achieved by adding *E. hellem* spores (30:1/spores: cells) to primary dendritic cells or DC2.4 cells cultures.

Successful invasion and colonization of *E. hellem* within cells or in host organs was verified by immuno-fluorescence assays and qPCR using *E. hellem* primers targeting at the conserved SSU-rDNA (5’-TGAGAAGTAAGATGTTTAGCA-3’; 5’-GTAAAAAGACTCTCACACTCA-3’) (5).

### S. aureus infection

For *in vivo* infection, the bacteria were inoculated into mice via intraperitoneal injection at the dose of 10^7^ CFU/mice. For *in vitro* infection, the bacteria were added to cell cultures at the ratio of 5:1 (DCs: *S. aureus*) or at the ratio of 1:1 (DCs: *S. aureus*), for various experimental purposes.

### Co-culture of DCs with T cells

DCs were infected with *E. hellem* (30:1/spores: cells) for 24 hours. The controls consisted of either un-infected cells or cells infected by *S. aureus* (5:1/DCs: *S. aureus*). These DCs were washed with PBS to get rid of excess pathogens, and then Jurkat cells were added to the culture groups (1:1 DCs/T cells). The innate immune cells and the lymphocytes were then co-incubated for 12 hours. After that, the flasks were gently stirred to collect cells in suspension (T-cells), while the cells gently stir the flasks to collect the suspension cells (T cells), the bottom-attached cells were collected as DCs, and were used for further analysis respectively.

### Flow cytometry analysis

Immune cell profiles and cell characterization, and the expressions of cell surface markers were all assessed by flow cytometry analysis. Single cell suspensions, from various treatments, were washed with 1x PBS/0.3 % BSA, and then stained with fluorochromes-conjugated antibodies (all purchased from BioLegend, USA) for 30 min at 4°C. Samples were then subjected to analysis via FACSCanto II flow cytometer (BD Biosciences, USA), and the data were analysed with FACSDiva software (v6.1.2).

### Cytokine expression analysis

Interleukin-6 (IL-6)_ and interleukin-12 (IL-12_ expression levels were detected by ELISA Kits (Thermo Fisher Scientific, USA); the samples analysed via ELISA were either plasma drawn from mice or cell culture supernatants. Mice peripheral blood was drawn with anti-coagulant sodium citrate added then the blood samples were centrifuged at 400x g for 10 minutes to isolate plasma from the supernatant.

### qPCR

To assess transcription levels of target genes, total RNAs of different cells such as DCs were extracted by TRIzol (Ambion, USA). The RNA samples were then reversely transcribed to cDNAs using High-Capacity RNA to cDNA Kit (Yeasen Biotechnology, China). qPCR assay was carried out according to Hieff® qPCR SYBR Green Master Mix instructions (Yeasen Biotechnology, China). The genes/primers information is shown in supplementary data **(S-Table 1)**.

### Label free quantitative mass spectrometry

DC2.4 cells, either un-infected or infected by *E. hellem*, were subjected to RIPA lysis buffer. The total protein samples from each group were then subjected to label free quantitative mass spectrometry (MS) analysis performed on a Q Exactive mass spectrometer (Thermo Scientific). The MS data were analyzed using MaxQuant software version 1.5.3.17 (Max Planck Institute of Biochemistry in Martinsried, Germany). The biological functions of proteins were annotated by Gene Ontology (GO) Annotation (Blast2GO, http://www.blast2go.com) and using the online Kyoto Encyclopedia of Genes and Genomes (KEGG) database (http://geneontology.org/). We screed for differentially expressed proteins between the control and *E. hellem*-infected groups by determining which proteins experienced a fold expression change greater than 1.5 fold (up to 1.5 fold or less than 0.5). These differentially expressed proteins were further analyzed usingseveral different bioinformatic analyses including hierarchical cluster by ComplexHeatmap R (version 3.4), KEGG Functional Enrichment analysis, and Protein-Protein Interact Network using IntAct molecular interaction database (http://www.ebi.ac.uk/intact/). Statistical significance was analyzed using Student’s t-test based on *P*-value < 0.05.

### Exogenous expression of *E. hellem*-PP1 in DCs

The *E. hellem* PP1 coding region was cloned into a vector with fluorescent tag (pCMV-mCherry-PP1). Transform the vector into isolated DCs using Lipofectamine 3000 reagent in Opti-MEM medium (Thermo Fisher Scientific, USA). The successful transformation and transient expression of E. hellem-PP1 was determined by tag fluorescence and immune-fluorescent assay.

### Statistics

Statistical analysis of results was conducted by using Student’s T-test or Two-way ANOVA and to identify the differences between two groups, with P < 0.05 being considered a significant difference.

## Results

### Persistent infection of *E. hellem* increases host disease susceptibility and disturbs dendritic cell populations

Microsporidia infections are usually asymptotic, however their covert infection and persistance within host may disturb immune responses, therefore making the host more vulnerable to further challenges. To test this hypothesis, we first utilized our previously established microsporidia infection model to infect wild-type C57BL/6 mice with *E. hellem*. The *E. hellem-infected* mice showed by no obvious symptoms such as spleen edema or significant weight loss (supplementary **S-Fig. 1A, 1B)**. However, *E. hellem* infection persisted as proved by detection of their spores in host blood, stool and urine samples even after more than 15 days post infection (supplementary **S-Fig. 1C**).

Interestingly, when *S. aureus* or endotoxin lipopolysaccharide (LPS) were inoculated into the *E. hellem* pre-infected mice, the hosts showed increased susceptibility to the secondary challenges. As shown, *E. hellem* pre-infection plus secondary infections (co-infection) caused significantly more weight loss and slower weight re-gain in mice, compared to secondary challenges alone (**Fig. 1A-1B**). Tissue and serum samples were collected and analyzed the endpoint of experiment. Results showed that *E. hellem* infection alone had no significant irritations to host organs such as spleen, but *E. hellem* pre-infection plus secondary infections (co-infection) led to more irritations in tissues compared to secondary challenges alone (**Fig. 1C-1D**). In addition, ELISA analysis of plasma cytokine levels revealed that *E. hellem* infection not only down-regulated the expression of pro-inflammatory cytokines IL-6 and IL-12, but also did not additively increase the cytokines levels when *E.hellem* infection along with the known cytokine stimulator LPS or *S. aureus* challenges (**Fig. 1E-1H**).

**Figure 1.**
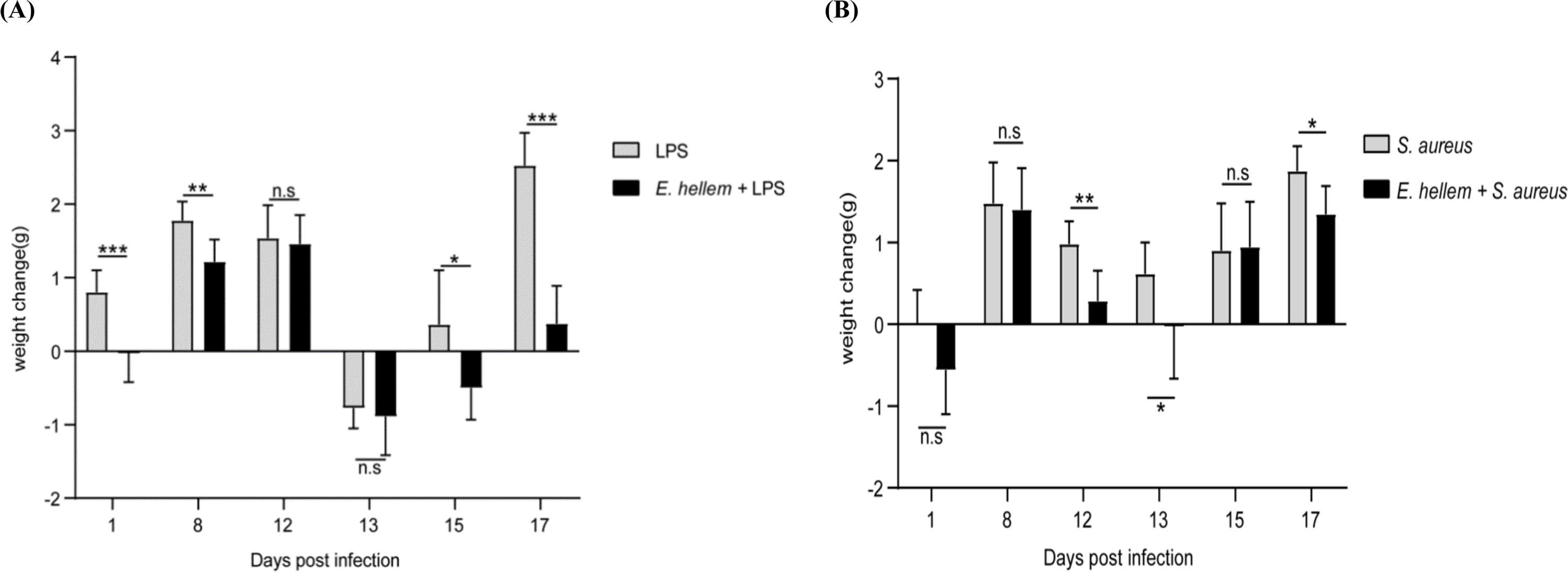
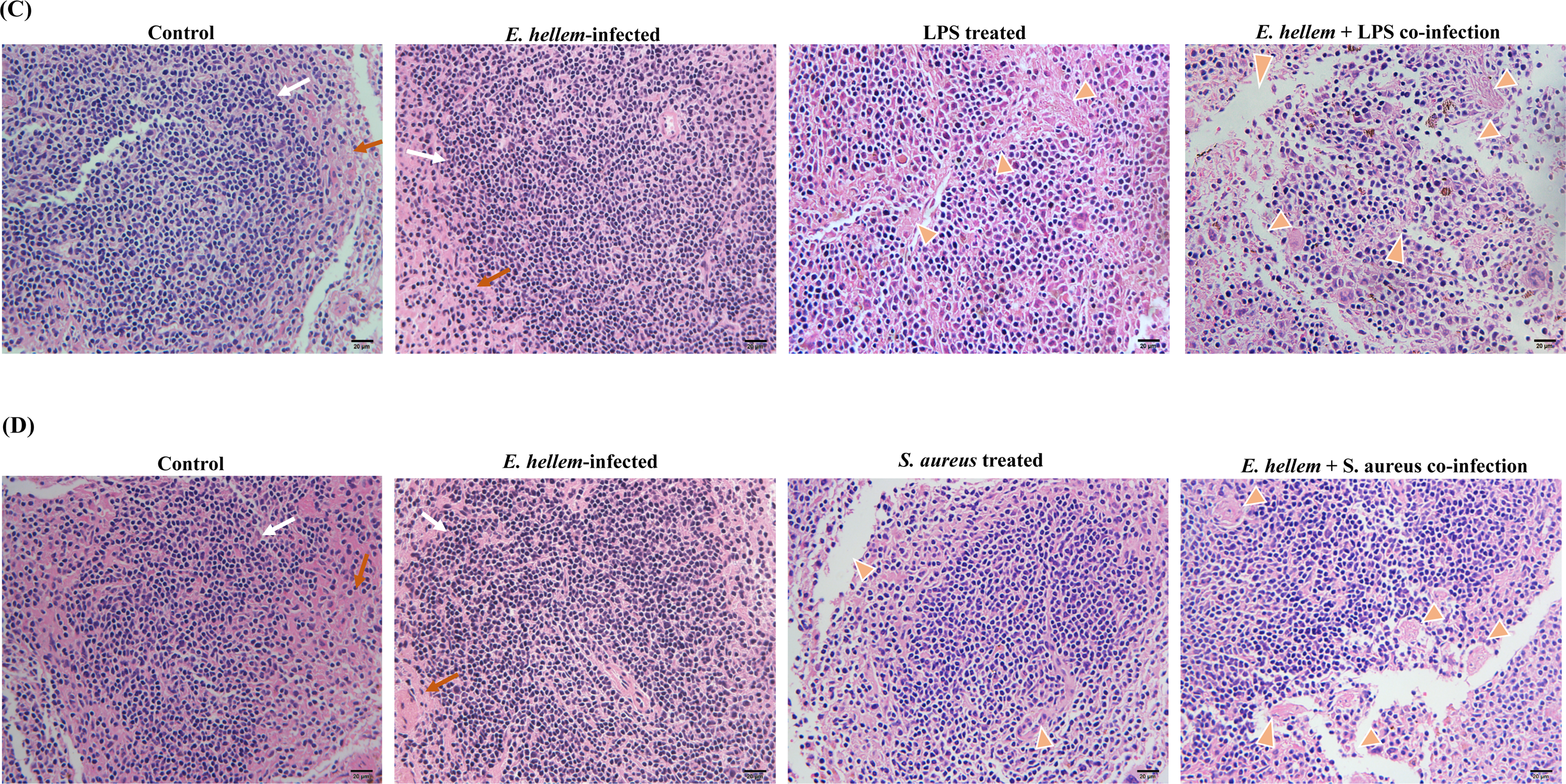
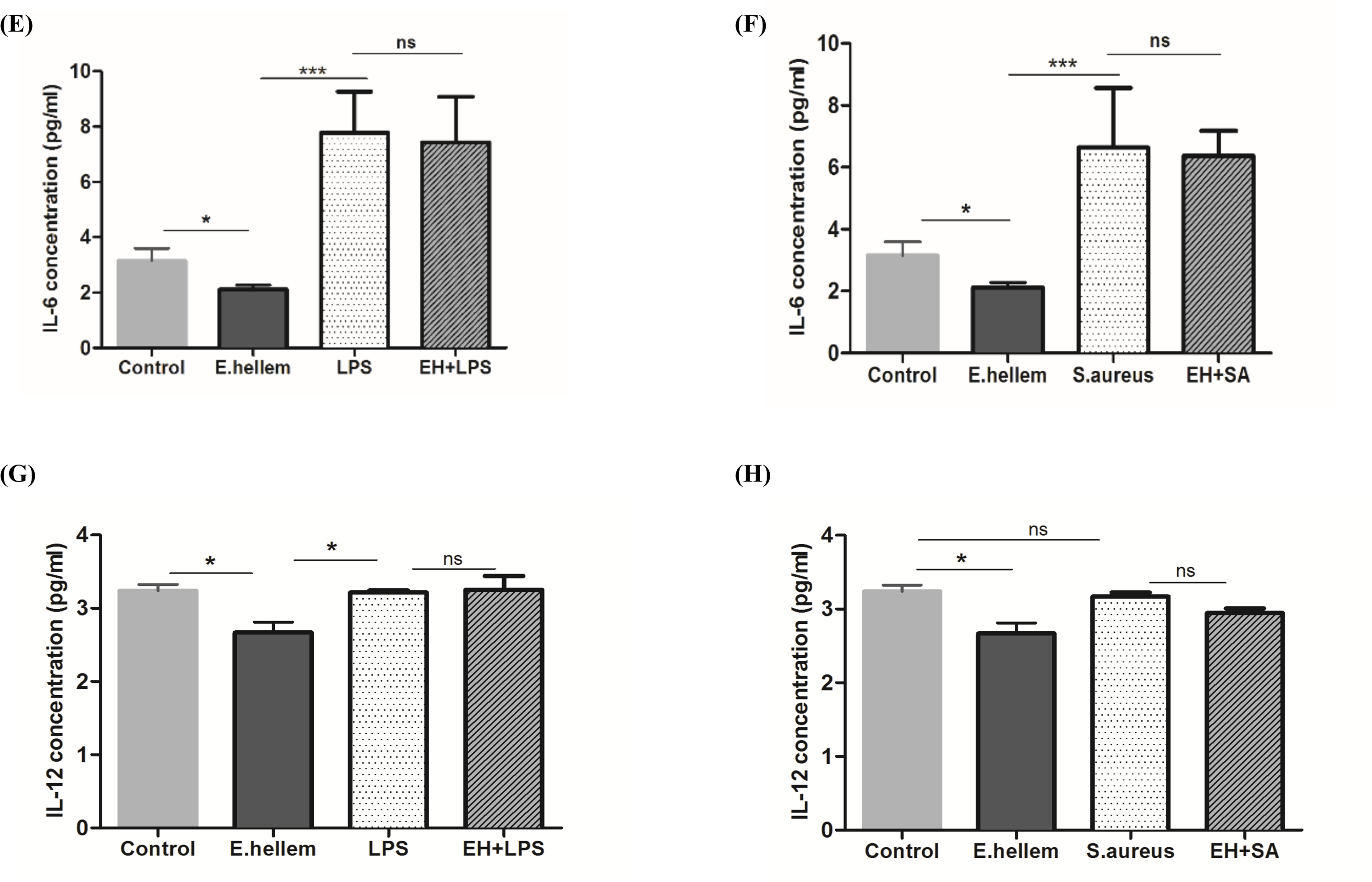
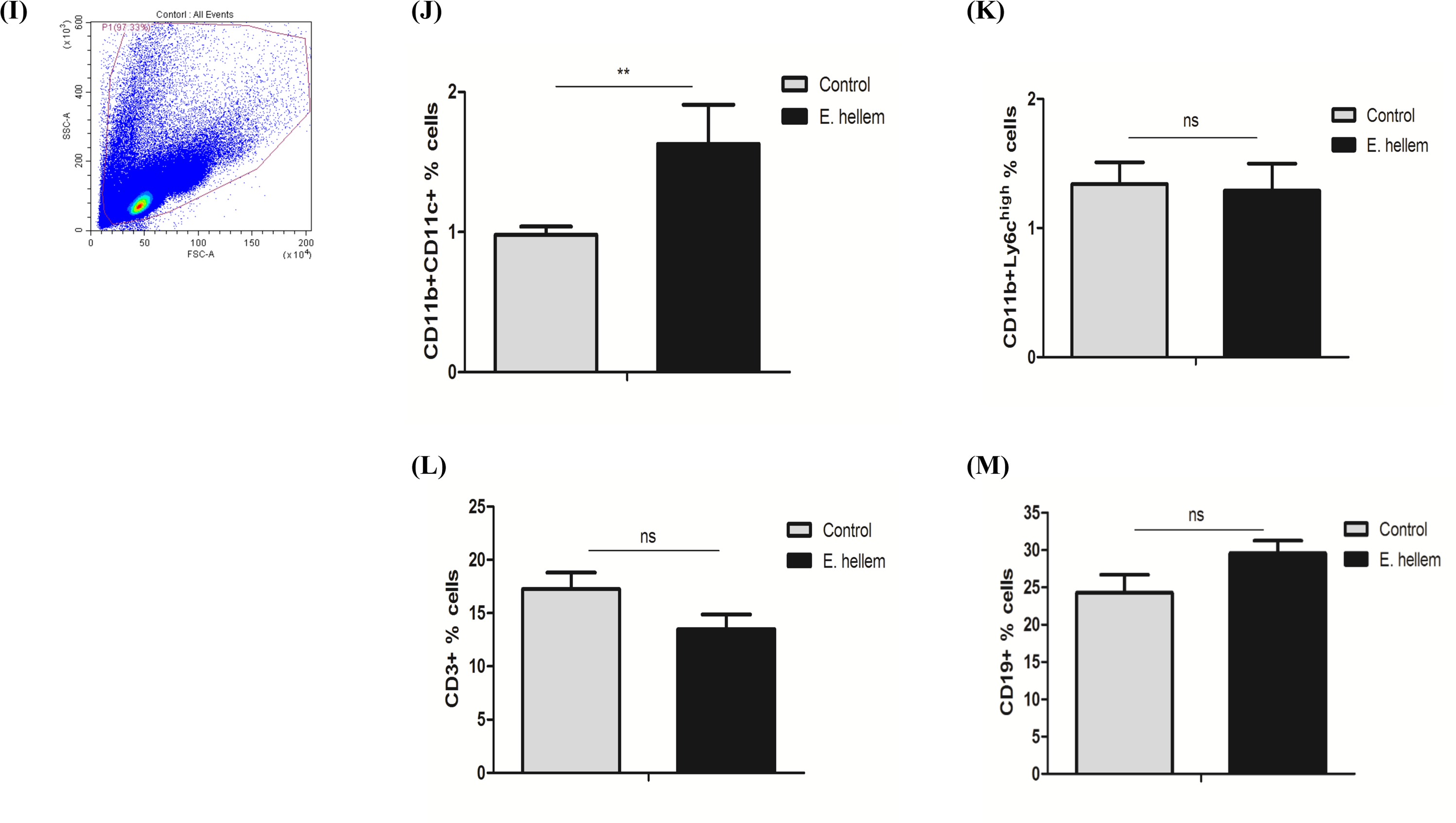
*E. hellem* infection and persistence increases host disease susceptibility and also affects dendritic cells. **(A, B)** LPS treatment or *S. aureus co-*infection in *E. hellem* pre-infected mice (*E. hellem*+*S. aureus*, *E. hellem*+LPS). Co-infections caused more body weight loss and slower/less body weight re-gain vs. single LPS or *S. aureus condition.* (n=8 per group) (*= P<0.05, **=P<0.01, ***=P<0.001, n.s. – not significant). **(C, D)** Hematoxylin-eosin staining of spleen samples from mice treated with either *E. hellem* or LPS or *S. aureus* alone, or with co-infections. *E. hellem.* Infection alone had no obvious effects on tissue pathology compared to control, as white pulps (white arrows) and red pulps (orange arrows) arranged normally. *E. hellem* pre-infection plus LPS treatment, or *E.hellem* pre-infection plus *S. aureus* infection, caused more damage to tissues compared to single challenges alone, as shown by more enlarged/distorted cells/cytosols and vacuolations (golden arrow heads) (Scale bar=20 ❍m). **(E-H)** ELISA assays to determine IL-6 (**E, F**) or IL-12 (**G-H**) levels in serum from mice with single *E.hellem* infections (E. hellem), LPS challenge (LPS), *S. aureus* (SA) and from *E.hellem* pre-infection plus LPS or *S. aureus* (EH+LPS/EH+SA). *E. hellem* infection significantly down-regulated the expression of both IL-6 and IL-12 compared to un-infected mice (Control); *E. hellem* pre-infection along with *S. aureus* infection (*EH+SA*) did not cause an additive increase of either IL-6 or IL-12, but a slight, albeit not-significant decrease of IL-12 levels were seen in EH+SA condition compared to *S. aureus* infection alone (**F, H**). A similar pattern was seen with LPS-treated mice; LPS single challenge significantly raised IL-6 and IL-12 levels above those seen in *E. hellem* infected mice, but *E.hellem* infection + LPS treatment (EH+LPS) did not additively increase IL-6 or IL-12 levels (**E, G**). (n=8 per group; *= P<0.05, **=P<0.01, ***=P<0.001, n.s – not significant). **(I-M)** Flow cytometry analysis profile of the immune cells isolated from mouse mesenteric lymph nodes (**I**). Mesenteric lymph nodes were isolated from control or *E. hellem* infected mice and were placed into a single cell mixture (n=3-5 lymph nodes from each mouse, 8 mice per group). Results showed that dendritic cell levels (CD11b+CD11c+; **J**) were significantly increased after *E. hellem* infection, but not inflammatory monocytes(CD11b+Ly6c_high_; **K**), T cells (CD3+; **L**), nor B cells (CD19+; **M**) were affected. **=P<0.01, ns=no significance; n=8/group.

Since the major infection route of microsporidia is through ingestion, the first cells to sample the invading pathogens are the phagocytes in the intestinal mucosa such as dendritic cells (DCs), which then are drained to mesenteric lymph nodes (MLN). Therefore, we were very interested to determine whether *E. hellem* infection dysregulated the immune cell populations in the MLN and which cell group was affected the most Flow cytometry analysis showed that only the dendritic cells (DCs) populations were significantly altered after infection, not the lymphatic T cells or B cells, or the inflammatory monocytes (**Fig. 1I-1J**). Taken together, these data demonstrated that *E. hellem* infection does persist within host, making them more vulnerable to further challenges, and that the most affected sentinel cell populations are DC.. Therefore, DCs may play key roles in microsporidia-host interactions.

### *E. hellem* interferes DCs immune functions and maturation

The major immune functions of DCs include pathogen phagocytosis, cytokine expressions, and antigen presentation. The abilities of DCs either from *E. hellem* infected mice or controls, to phagocytose fluorescently-labelled *S. aureus* were compared and analysed. As shown, DCs from *E. hellem*-infected mice were clearly deficient in engulfment of *S. aureus*as compared to DCs from uninfected mice **(Fig. 2A**, and supplementary videos **S-Video1-4)**.

**Figure 2.**
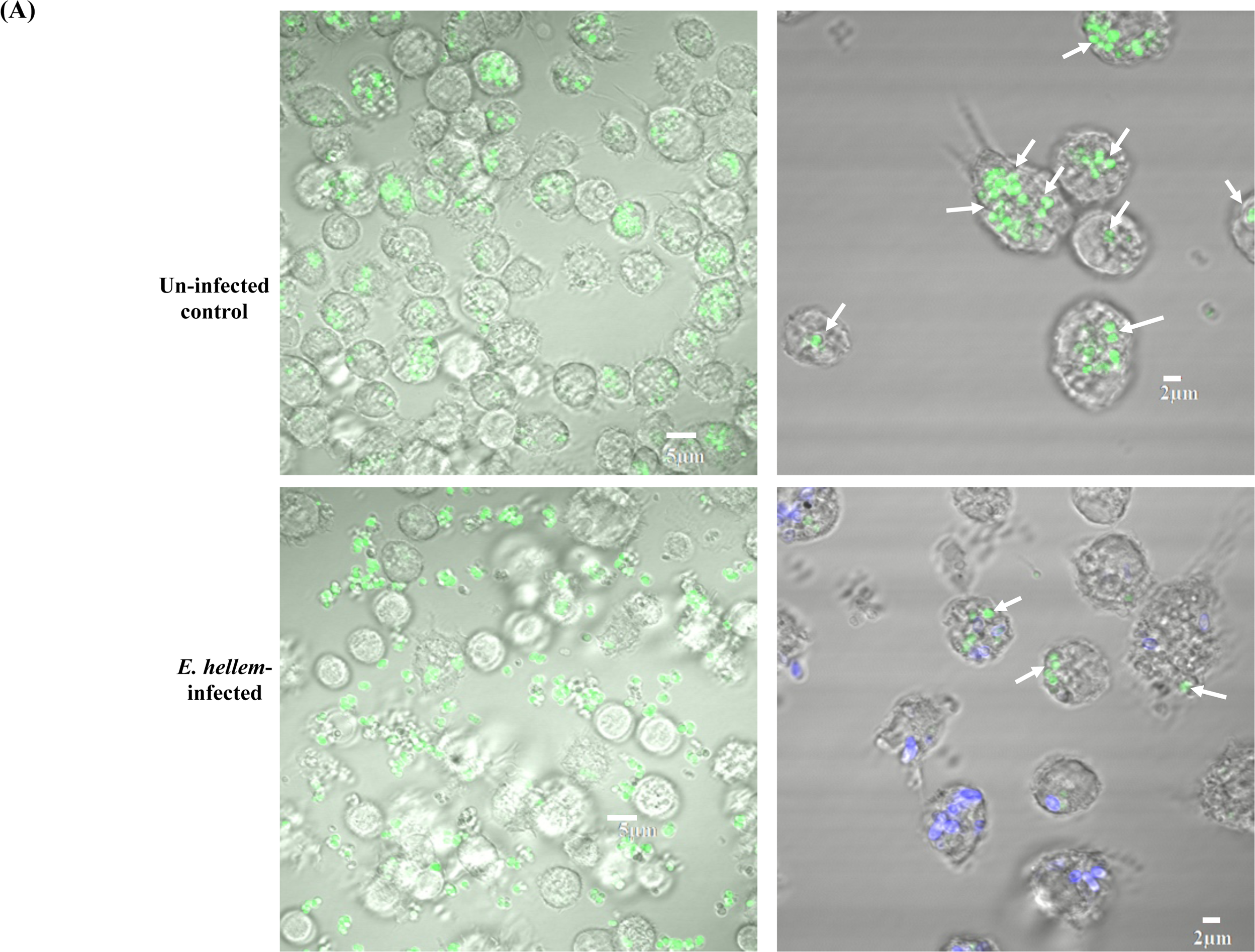
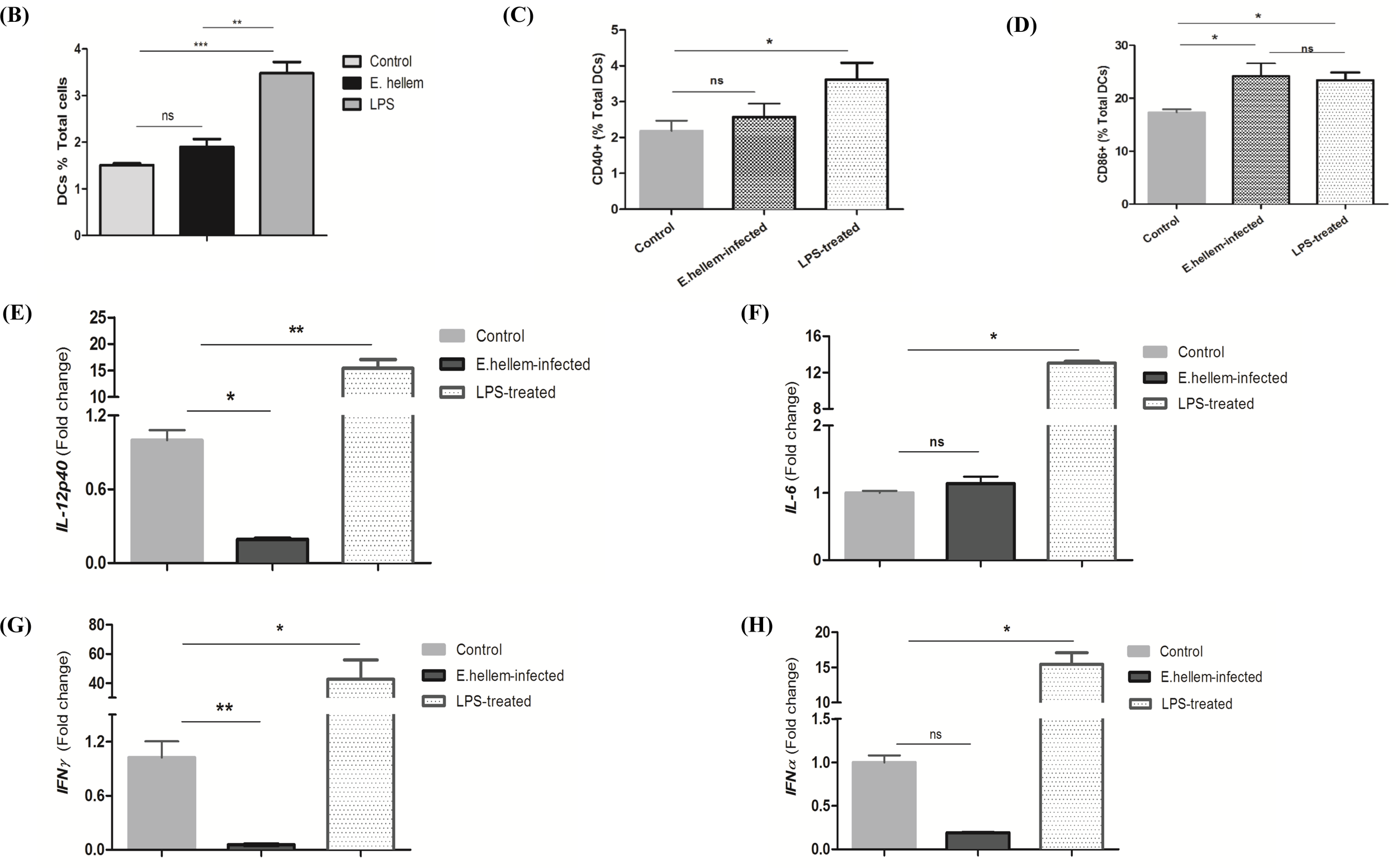
*E. hellem* interferes with the immune functions and maturation of DCs. **(A)** DCs were isolated from un-infected control mice or *E. hellem* infected (*E. hellem*-infected) mice, then fluorescent (GFP) labelled *S. aureus were* added to DCs culture (MOI=20:1). The phagocytic ability of the different DC populations was assessed by fluorescent microscopy. Most of the added *S. aureus* (green) were engulfed by control DCs, while most added *S. aureus* (green) were found floating outside of DCs isolated from *E. hellem*-mice (scale bar=5 ❍m). Magnification of images showed that DCs from *E. hellem*-infected group engulfed significantly less *S. aureus* (green; arrows) compared to DCs from un-infected controls. Persistence of *E. hellem* spores was manifested by calcofluor white stain (blue) (scale bar=2 ❍m). **(B)** Flow cytometry analysis showed that *E. hellem* infection suppressed the levels of the splenic dendritic cells (33D1+) to the comparable level as the un-infected control, but LPS treatment significantly up-regulated the dendritic cells. (n=8/group; ns=no significance, **=P<0.01, ***=P<0.001). **(C-D)** *E. hellem* infection, but not LPS treatment, inhibited the expressions of CD40 on dendritic cells, while the expression of CD86 on dendritic cells increased with both *E. hellem* infection and LPS treatment (n=8/group; ns=no significance, *=P<0.05). **(E-H)** Cytokine expression levels from splenic dendritic cells (DCs) isolated from control, *E.hellem* infected, or LPS-treated mice. **(E)** *IL-12p40* expression level of DCs from *E. hellem* infected mice were significantly down-regulated compared to un-infected mice and LPS-treated mice. **(F)** *IL-6* expression levels of DCs from *E. hellem* infected mice was unchanged from un-infected controls and significantly lower than from DCs isolated from LPS-treated mice. **(G)***IFN*_γ_ expression levels of DCs from *E. hellem* infected mice were significantly down-regulated compared to un-infected controls and LPS-treated mice. **(H)** *IFN*_α_ expression levels of DCs from *E. hellem* infected mice had no significant change between un-infected controls and were significantly lower than the LPS-treated ones. n=8/group; ns=no significance, *=P<0.05, **=P<0.01, ***=P<0.001.

Full maturation and efficient homing of DCs from MLN to T cell-region constitute crucial aspects of DCs functions. Therefore, we collected immune cells from the spleens of either *E. hellem*-infected, LPS-treated or un-infected mice to determine if *E.hellem* infection caused a defect in DC homing. Flow cytometry analysis demonstrated thatthe matured and specialized DCs populations were retained at comparable levels to un-infected controls even after *E. hellem* infection, but the LPS-treatment significantly increased the population of matured DCs in spleen (**Fig. 2B**). Flow cytometry analysis further confirmed that *E. hellem* suppressed the expansion of maturation and co-stimulatory surface markers, CD40 and CD86, positive DCs (**Fig. 2C-2D**). Next, the cytokine expression of splenic DCs were assessed by qPCR analysis and we confirmed that DCs from the *E. hellem* infected group expressed significantly lower levels of *IL-12p40*, *IFNγ* and *IL-6 IFNα,* as compared to DCs from un-infected controls and LPS treated groups (**Fig. 2E-2H**).

Taken together, these findings demonstrated that *E. hellem* infection interferes with the full immune functions and maturation processes of DCs, which would explain the above data which showed lower cytokine levels in serum and increased host susceptibility to secondary pathogens.

### *E. hellem* detained DCs antigen presentation and T cell priming potencies

Fully functioning DCs present the processed antigens to T cells and prime T cell activation. Therefore, we analyzed the expressions of *H2Aa* and *H2Ab,* in the MHC-II complex, along with *DC-SIGN*, all are essential for antigen presentation and T cell priming (39). qPCR assay analysis showed that, although *E. hellem* infection caused the up-regulation of *H2Ab*, it suppressed the up-regulations of *H2Aa* in the MHC-II complex and the essential antigen presentation marker *DC-SIGN* (**Fig. 3A-3C**); this data indicated that the antigen presentation abilities of *E. hellem* infected-dendritic cells would be impaired. Next, we analysed the alterations and stimulation markers of T cells populations isolated from the spleens of control and *E. hellem* infected mice by flow cytometry. Our results showed that neither the CD4+ T cells nor CD8+ T cells showed significant stimulation after *E. hellem* infection (**Fig. 3D-3E**). qPCR analysis of the expression of *Ctla4* and *Tigit*, the known markers for T cell activation, confirmed that there were no significant changes after *E. hellem* infection (**Fig. 3F-3G**). These data demonstrated that *E. hellem* supressed the antigen presentation abilities of DCs and T-cell priming potencies, as reflected by no up-regulation of MHC-II complex or DC-SIGN, and changes in stimulation markers of T-cell populations.

**Figure 3.**
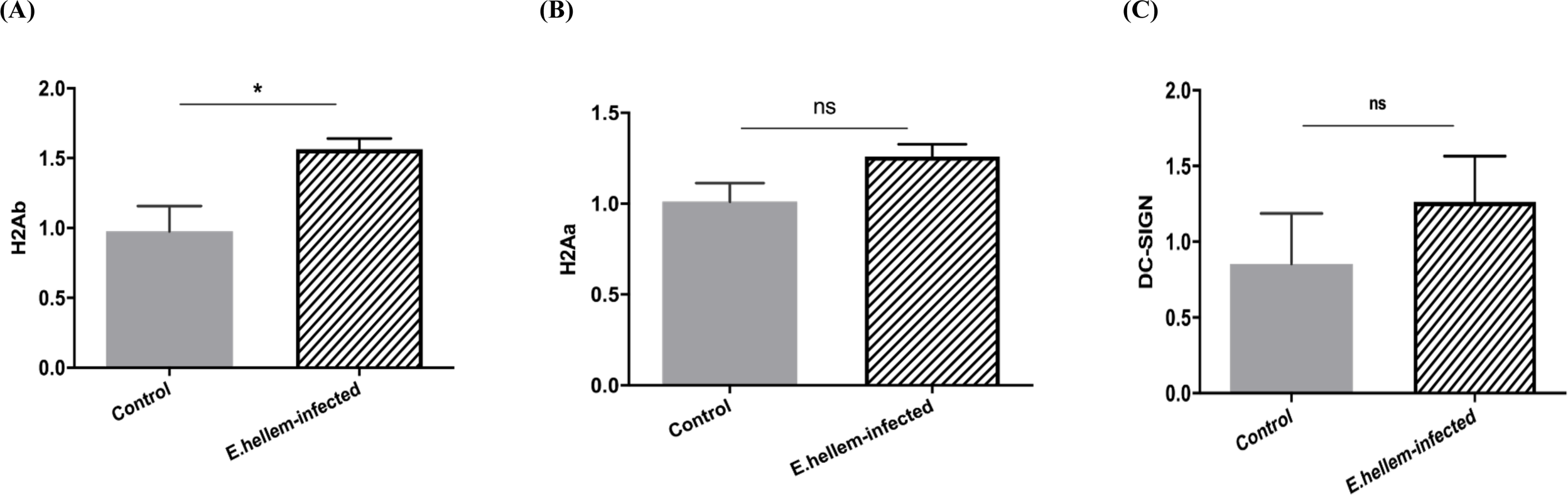
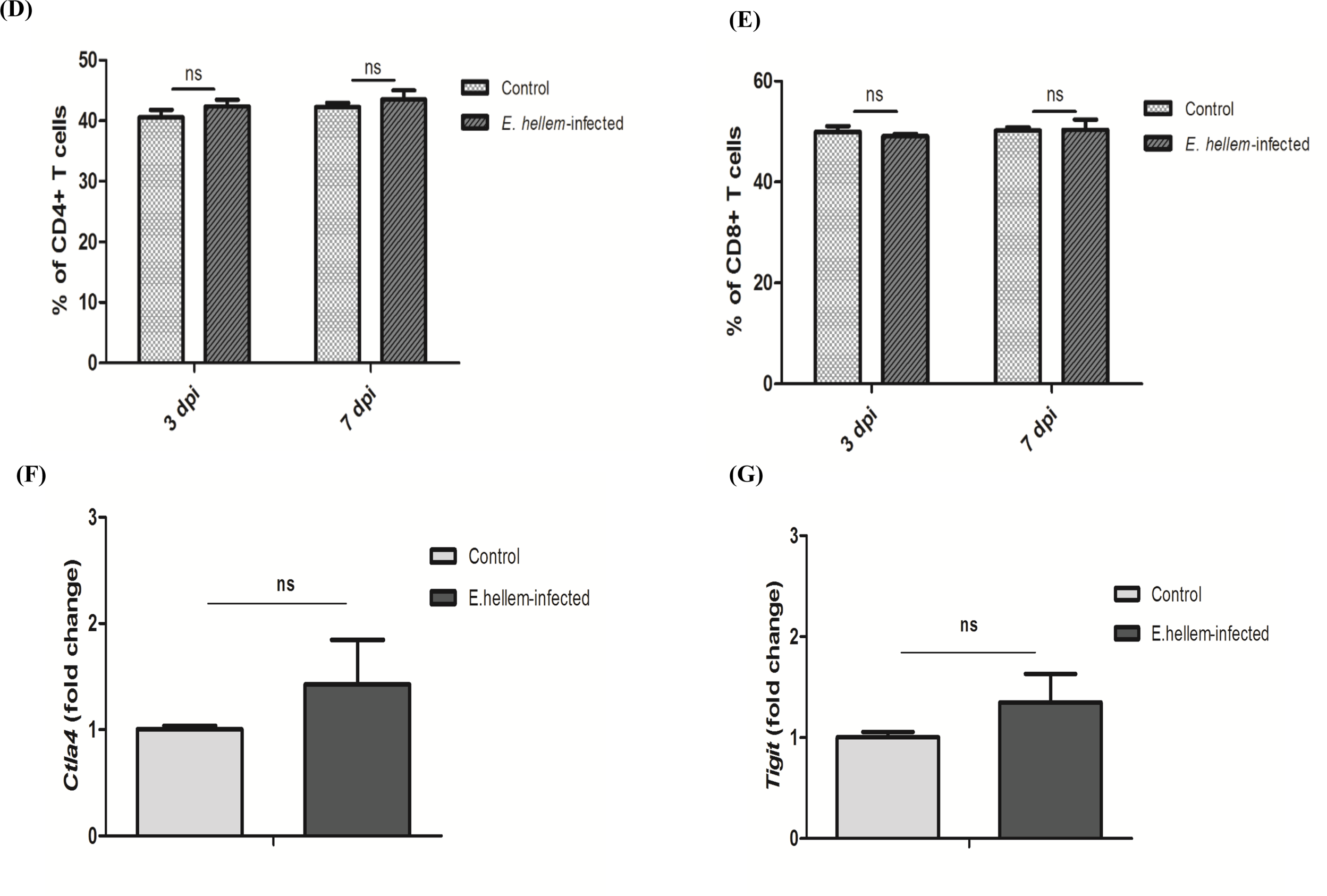
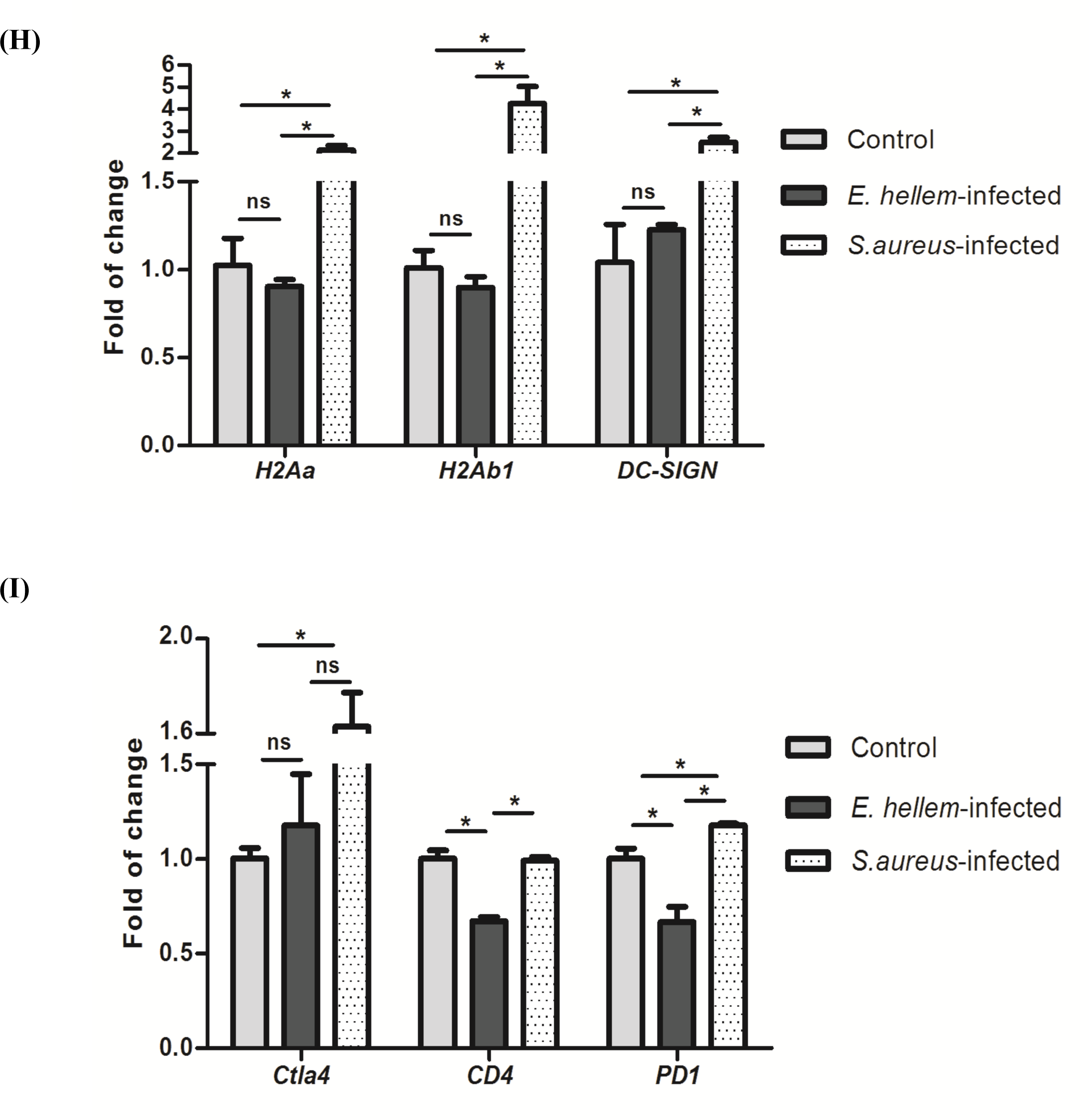
*E. hellem* suppresses the antigen presentation abilities and T cell priming potencies of DCs. **(A-C)** The expressions of DCs antigen-presentation related surface markers were assessed by qPCR assay. *E. hellem* infection caused the up-regulation of *H2Ab* marker (**A**), but suppressed the up-regulation of both *H2Aa* (**B**) and *DC-SIGN* (**C**)*. (*ns=no significance; *=P<0.05). **(D-E)** Levels of CD4+ T cells and CD8+ T cells were assessed after *E. hellem* infection at both 3 and 7 days post infection (3 dpi and 7 dpi). No population changes were detected after *E. hellem* infection. (n=8/group; ns=no significance). **(F-G)** The expressions of *Ctla4* and *Tigit in* T-cells after *E.hellem* infection were analysed by qPCR analysis. No statistical significance in the expression of these activation matters between *E.hellem* infected and un-infected control cells. (ns=no significance). (**H**) DC2.4 cells were first infected by *E. hellem*, *S. aureus* or uninfected (control) and then were co-cultured with Jurkat-T cells; the cells were separated later for surface marker analysis. qPCR results showed that the expressions of antigen-presentation markers tested, *H2Aa*, *H2Ab1*, and *DC-SIGN*, showed down-ward trends by *E. hellem* infection, compared to *S. aureus* infections. (ns=no significance; *=P<0.05). **(I)** The Jurkat T-cells isolated from co-culture with DC2.4 were also analyzed by qPCR. Expressions of representative markers, *Ctla4, CD4 and PD-1,* also showed down-ward trends after *E. hellem* infection, with statistically significant down-regulation of *CD4* and *PD-1*(n=10/group; ns=no significance; *=P<0.05).

Next, we took another step forward by co-culturing DC2.4 cells and Jurkat-T cells together *in vitro,* to further demonstrate the interference of *E. hellem* on DCs functions and reluctance of T-cell priming. DC2.4 cells were infected by *E. hellem* or *S. aureus* and then co-cultured with Jurkat-T cells *in vitro* for 12 hours. DCs and T-cells were then separated and their total RNAs were extracted for qPCR analysis. Our results again confirmed that the expressions of *H2Aa*, *H2Ab* and *DC-SIGN* of *E. hellem*-infected DCs showed down-ward trends compared to *S. aureus* infected DCs (**Fig. 3H**); the expressions of T-cell activation markers such as *CD4*, *Ctla4*, and *PD-1* also showed down-ward trends or even statistically significant down-regulated by *E. hellem* infection (**Fig. 3I**).

Taken together, our findings demonstrated that *E. hellem* infection supresses immune functions of DCs and T-cell priming potencies. We surmise that the impaired innate immune responses by of *E. hellem* infected DCs coupled together with the subsequent suppression of T-cell stimulation would severely weaken host immunity against any further co-infections or challenges.

### The MAPK-NFAT5 signaling pathway is key for *E. hellem* -DCs interaction and modulations

We have demonstrated that *E. hellem* down-regulated the expression of several immune related genes such as *IL-6*, *H2Ab* and *H2Aa* in DCs. It is known that in immune cells, the expressions of these genes were often regulated by transcriptional factor NFAT5, often activated by p38α-MAPK signaling cascade. Therefore, we investigated the proteome changes of DCs after *E. hellem* infection by mass spectrometry (Top differentiated DCs proteins were listed in supplementary **S-Table 2)**. Our sub-cellular localization analysis revealed that most of the differentially expressed proteins of DCs were localized in the cytoplasm or nucleus (**Fig 4A**), indicating that many interactions and modulations induced by *E. hellem* occurred in cytoplasm or nucleus. Gene Ontology (GO) enrichment as well as KEGG analysis showed that E. hellem infection affected many cellular events in DCs, such as signaling pathways including the MAPK signaling pathways/cascade. (**Fig. 4B**). We list the top differentially expressed DC proteins, particularly ones associated with cellular responses, signal transduction, and transportation in **Table 1**. The protein-protein interactions (PPI) network analysis of these top identified proteins revealed that MAPK signaling pathway isof the central links of this representative PPI network (**Fig. 4C**).

**Figure 4.**
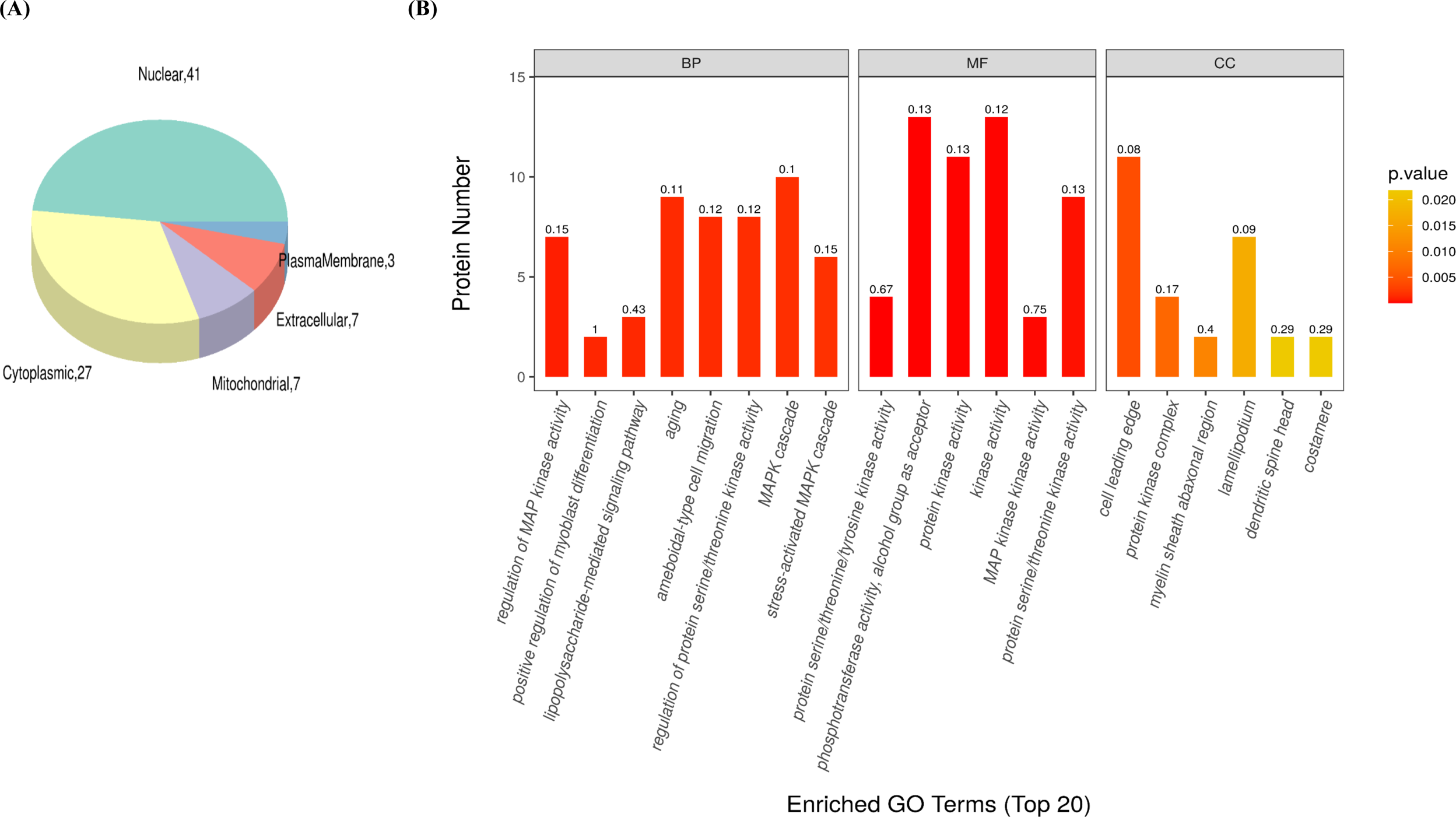
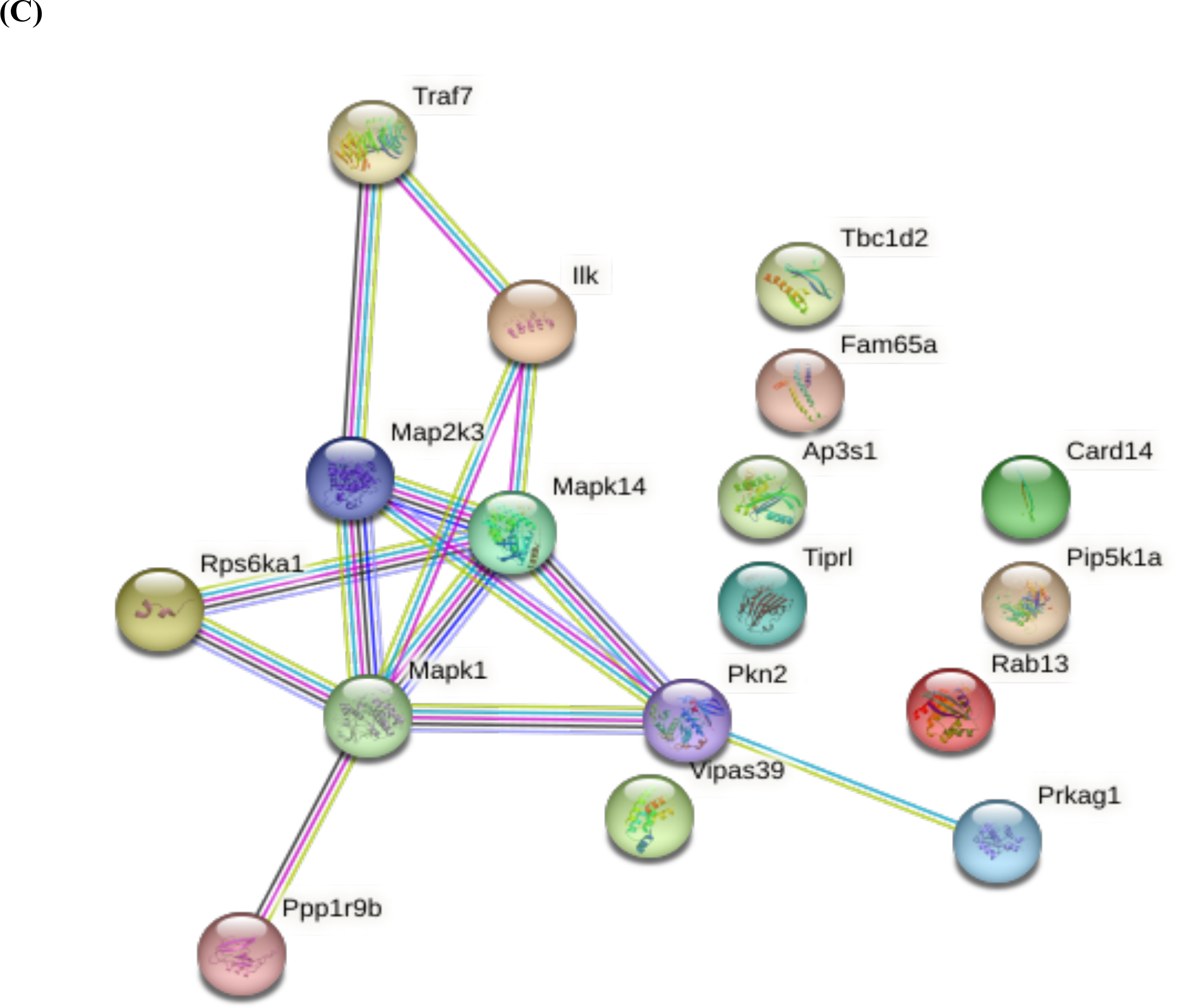
p38α /MAPK signaling pathway is key for *E. hellem* -DCs interaction and modulations. **(A)** Pie chart of sub-cellular localizations of the most differentially expressed proteins in DCs after *E. hellem* infection vs control DCs; expression of 41 nuclear and 27 cytoplasmic proteins are significantly altered during *E. hellem* infection. **(B)** GO enrichment analysis of the top differentially expressed DCs proteins after *E. hellem* infection. Multiple signaling pathways, including the MAPK pathway, were among the most affected cellular events. **(C)** Protein-protein interaction network, showing several of the top differentially expressed DCs proteins during *E. hellem* infection were associated with MAPK signaling pathway.

**Table 1.**
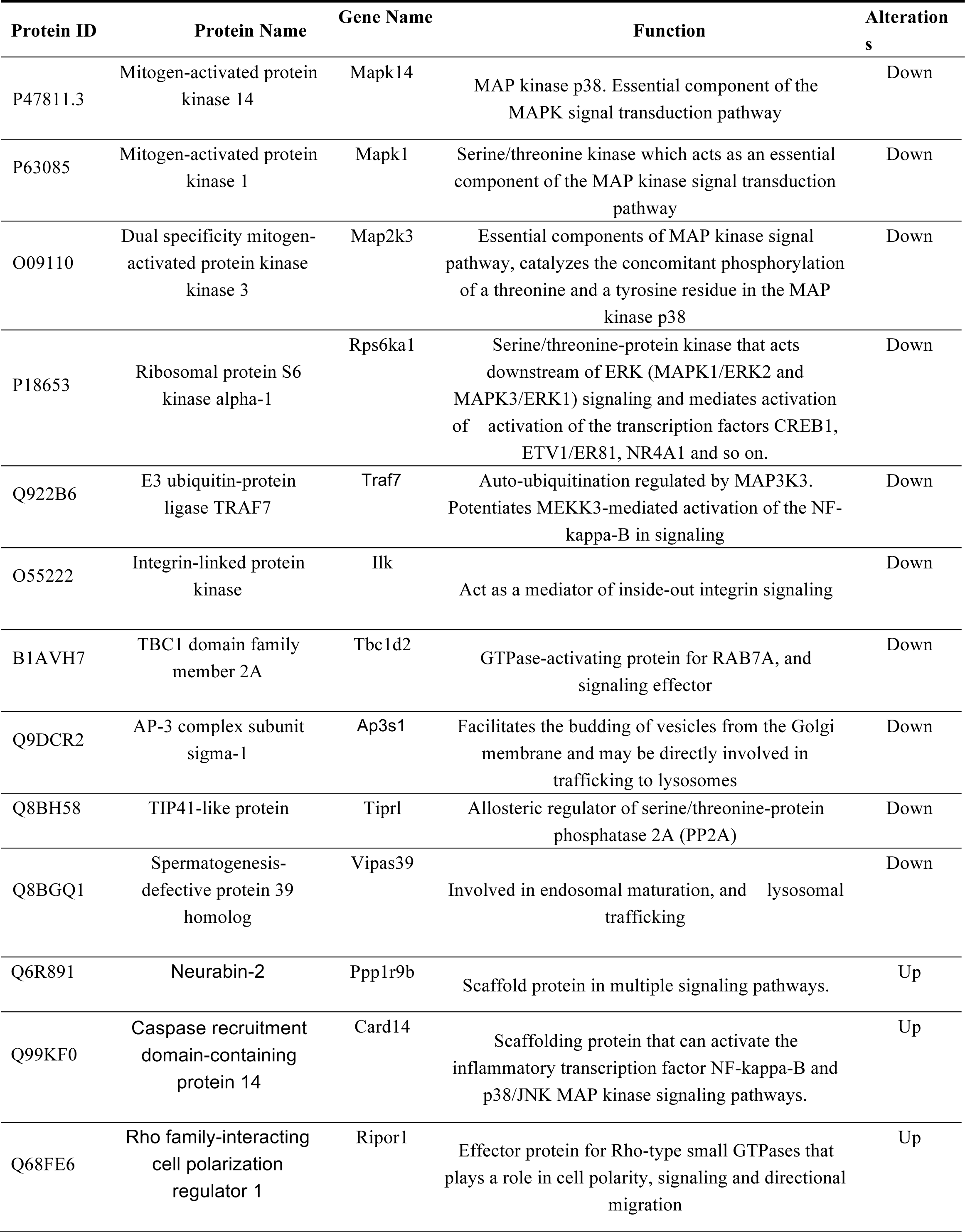
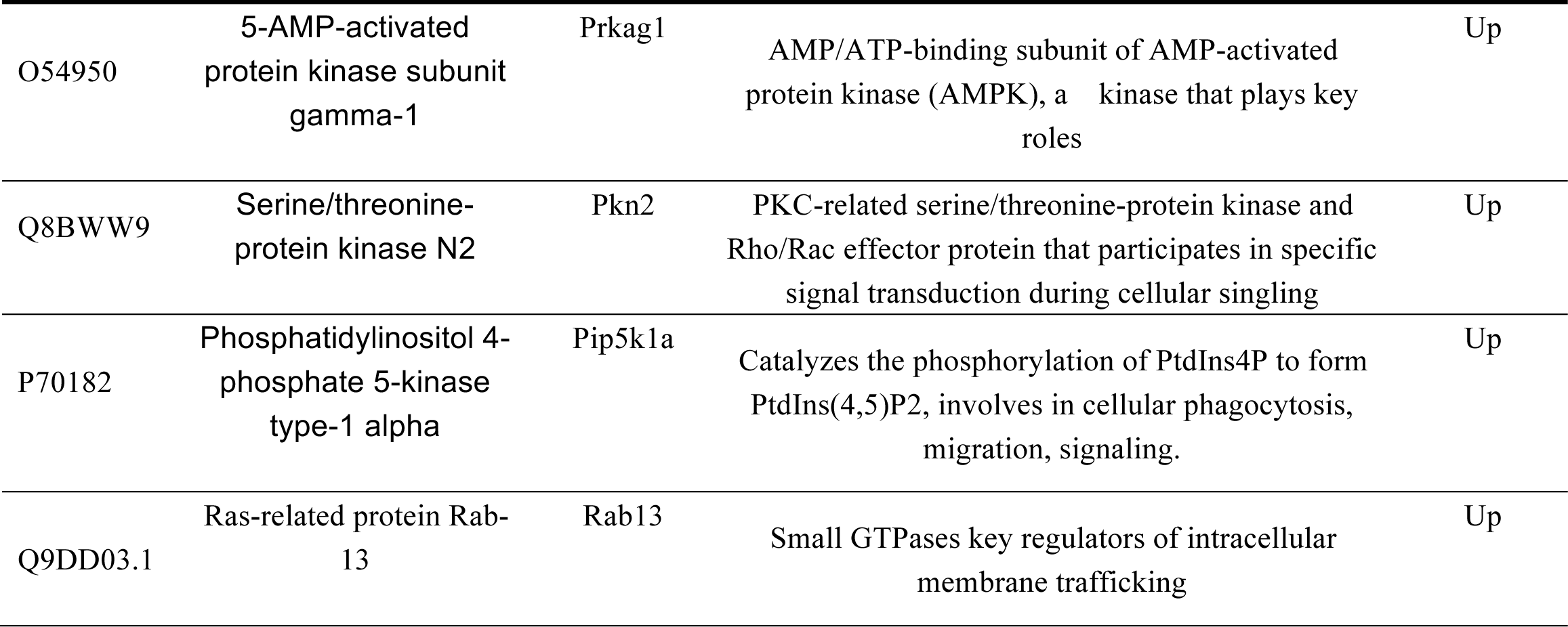
Representative DCs proteins differentially expressed after *E. hellem* infection.

Next, we assessed the expression and localization of NFAT5 after *E. hellem* infection. Western blot assay demonstrated that the NFAT5 expression was suppressed by *E. hellem* infection in DC2.4 cells, compared to un-infected and S. aureus infection controls (**Fig. 5A**). Immuno-fluorescent assay revealed the translocation of transcriptional factor NFAT5 from cytoplasm into nucleus was severely inhibited by *E. hellem* infection (**Fig. 5B**). To verify the essential roles of NFAT5 in the response of DCs g to *E. hellem*, we knocked-down NFAT5 in DC2.4 cells by RNAi (Fig. 5C) and demonstrated that *E. hellem* proliferation increased almost 2-folds (**Fig. 5D**). These findings suggested that NFAT5 and the related MAPK signaling pathway are a key axis in responding to *E. hellem* infection, therefore may be the major regulation targets of pathogen-host interactions.

**Figure 5.**
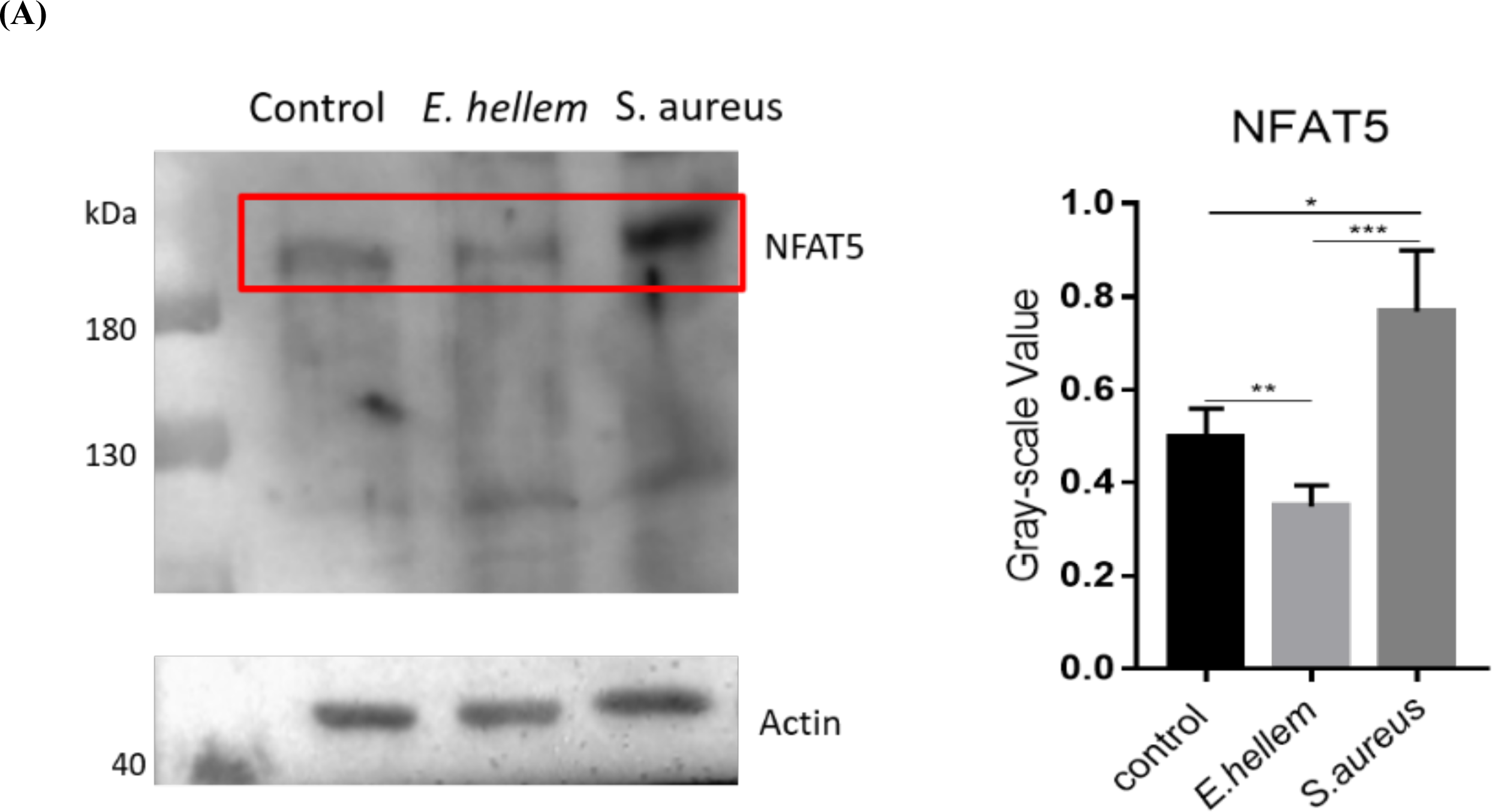
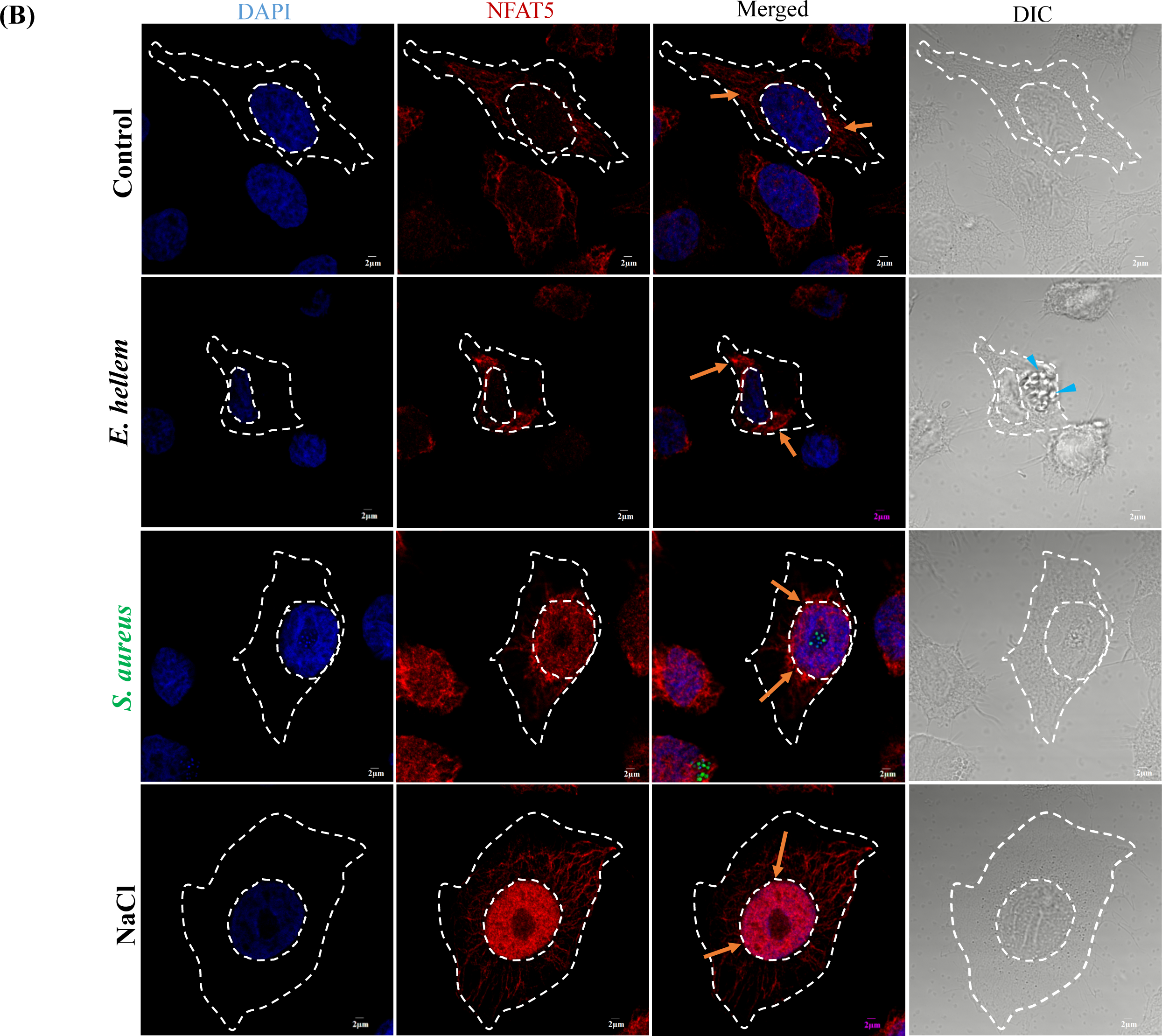
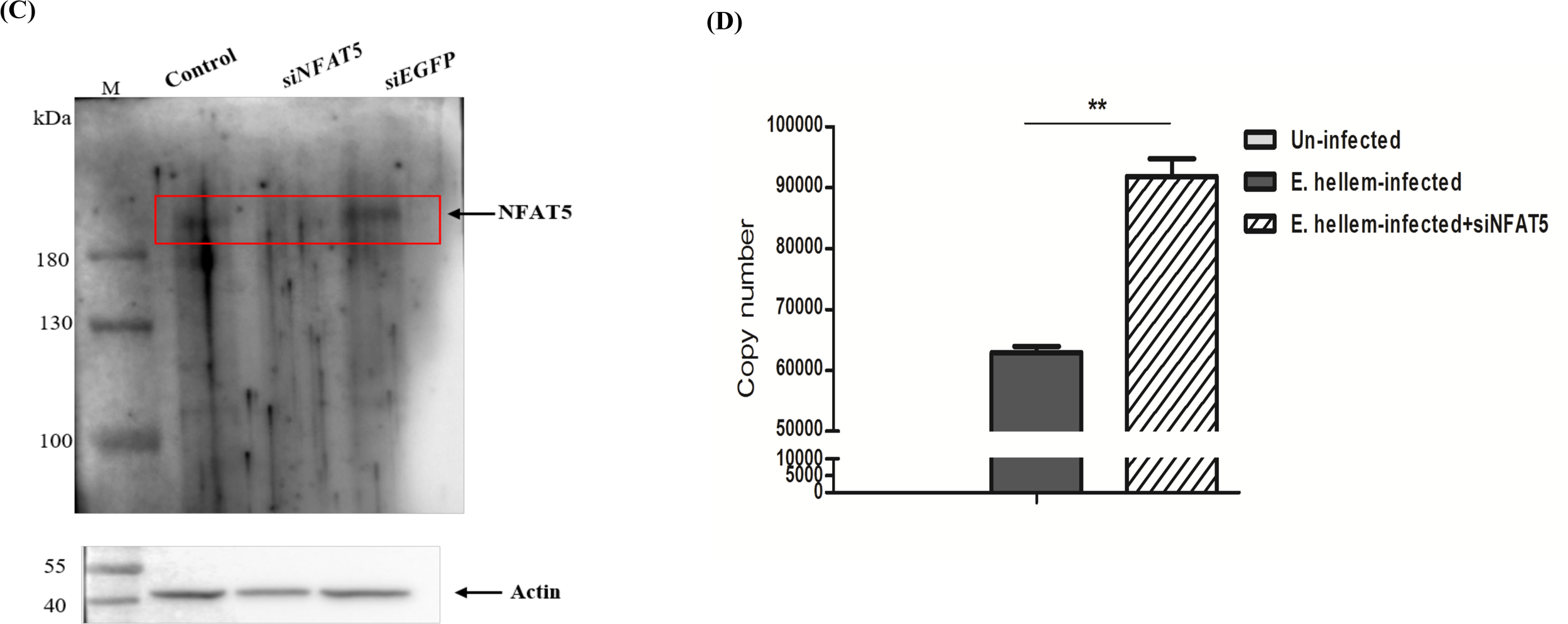
NFAT5/MAPK signal axis is essential during *E. hellem* infection and cellular modulation. **(A)** Western blot analysis of NFAT5 protein levels in DC2.4 cells under un-infected conditions (control), infected with *E. hellem* or with *S. aureus*. NFAT5 expression was suppressed by *E. hellem* infection, compared to control or *S. aureus* infection conditions. NFAT5 Western blots were analyzed via densitometry and gray scale values of NFAT5 bands under aforementioned test conditions were charted. (*=P<0.05, **=P<0.01, ***=P<0.001). **(B)** Immunofluorescent microscopy of NFAT5 localization in DCs under control conditions or under either *E. hellem* or *S. aureus* infection. NFAT5 (Alex Flour 594) is constitutively expressed in the cytoplasm (orange arrows) in control DCs and NFAT5 localization remained in the cytoplasm (orange arrows) after *E. hellem* infection. *E. hellem* was able to proliferate in DCs and form parasitophorous vacuoles (blue arrows). Both *S. aureus* infection and osmotic stimulation via salt (NaCl) lead to up-regulation of NFAT5 expression and nuclear re-localization (orange arrows) (scale bar=2 ❍m). **(C)** Western blot analysis of NFAT5 protein levels in DC2.4 cells after RNAi treatment with NFAT5 specific RNAi (siNFAT5) or nonspecific RNAi (siEGFP); NFAT5 was significantly down-regulated after RNAi interference with siNFAT5 vs siEGFP and control conditions. **(D)** qPCR analysis of *E. hellem* proliferation within host DC2.4 cells, with and without siNFAT5 treatment. Disruption of NFAT5 via siNFAT5 significantly increased *E. hellem* proliferation, as reflected by the copy numbers of *E.hellem* specific *PTP4* (n=8/group; **=P<0.01).

### *E. hellem* Serine/Threonine Protein Phosphatase (PP1) targets DCs p38α (MAPK14)/MAPK

To identify the regulating factors from *E. hellem*, we analyzed *E. hellem*-derived proteins within infected DCs by mass spectrometry (*E. hellem* proteins identified in DCs were listed in supplementary **S-Table 3**). Results revealed that most of these proteins are associated with protein binding, cellular responses and signal transduction. Moreover, we found that the serine/threonine protein phosphatase PP1 is one of the top expressed *E. hellem*-derived proteins in DCs (Representatively identified *E. hellem* derived proteins in **Table 2**).

**Table 2.** Representative *E. hellem* proteins identified in infected DCs.

| **Protein ID** | **Gene ID** | **Protein Name/Function** |
| --- | --- | --- |
| XP_003888309.1 | 13466767 | PP1 serine/threonine phosphatase |
| XP_003886772.1 | 13467540 | Ras-like GTP binding protein |
| XP_003887182.1 | 13468215 | GTP-binding nuclear protein |
| XP_003887892.1 | 13466917 | Rab GTPase |
| XP_003886950.1 | 13467595 | Beta-tubulin |
| XP_003886661.1 | 13466496 | Eukaryotic translation initiation factor 2 subunit gamma |
| XP_003886837.1 | 13467379 | RAD3-like DNA-binding helicase |
| XP_003887105.1 | 13467910 | Nop56p-like protein |
| XP_003887061.1 | 13468065 | DNA topoisomerase II |
| XP_003886739.1 | 13467235 | Dihydrofolate reductase |
| XP_003887411.1 | 13467332 | Histidyl-tRNA synthetase |
| XP_003887230.1 | 13468325 | Hypothetical protein |
| XP_003887352.1 | 13467170 | Hypothetical protein |
| XP_003887011.1 | 13467738 | Hypothetical protein |
| XP_003887459.1 | 13467461 | Hypothetical protein |

PP1 is a major Ser/Thr phosphatase, highly conserved in all eukaryotes and is known to regulate p38α/MAPK signaling at several levels, along with directly interacting with component proteins in the pathway. We firstly analyzed the sequence homology of *E. hellem*-PP1 with mouse and human PP1s (**Fig. 6A**). Result showed the *E. hellem*-PP1 is highly conserved and possess the potential to mimic the binding/regulating functions of host PP1 on signaling pathways. To verify the direct interactions of *E. hellem*-PP1 with host proteases, we next utilized yeast two hybrid assay, and confirmed that *E. hellem*-derived PP1 directly interacts with DCs P38α (MAPK14) (**Fig. 6B**). Moreover, we expressed *E. hellem*-derived PP1 in normal DC cells (**Fig.7A**), and showed that the ectopically-expressed *E. hellem* PP1 co-localized with host p38α (MAPK14) (**Fig.7B**). In addition, the expression of representative genes from the MAPK pathway such as NFAT5, IL-6, H2Aa and H2Ab were all significantly down-regulated by ectopically expressed *E. hellem* PP1 (**Fig.7C**).

**Figure 6.**
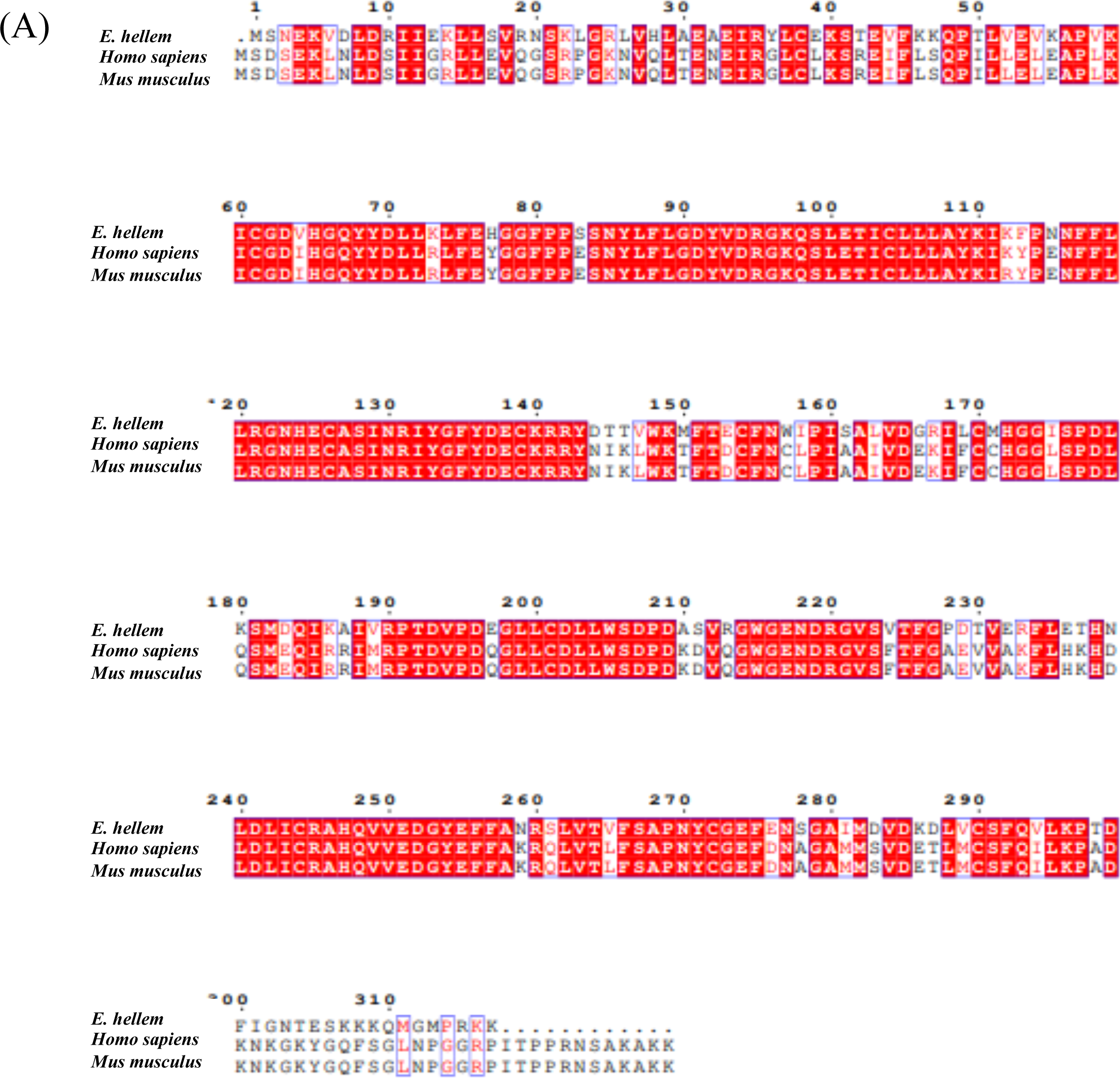
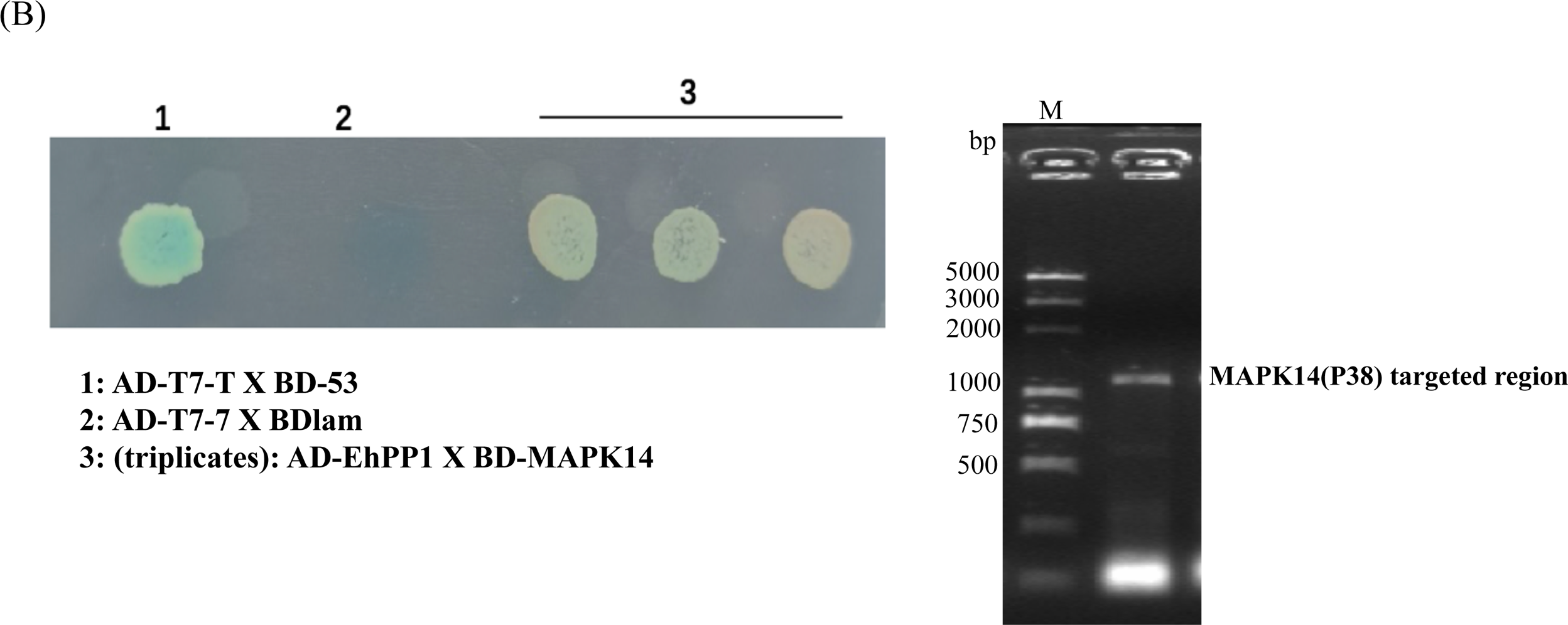
Direct binding of *E. hellem*-PP1 with DC-p38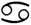(MAPK14), shown by yeast two-hybrid assay. **(A)** Protein sequences alignment of Ser/Thr protein phosphatase (PP1) from *E. hellem*, mouse and human, showed high conservations among the proteins. **(B)** MAPK14 expressed by DCs was cloned into pGBK7 plasmid (BD-MAPK14), and *E. hellem*-Serine/Threonine Protein Phosphatase PP1 was cloned into pGADT7 plasmid (AD-EhPP1). These plasmids were then transformed into competent yeast cells and the binding between MAPK14 and PP1 was validated in synthetic dropout-Leu-Trp -Ade-His medium supplemented with Xα-gal. The fusion strain of pGBKT7-53 with pGADT7-T was used as the positive control, fusion strain of pGBKT7-lam with pGADT7-T was used as the negative control. The EhPP1 and MAPK14 fused clones were subjected to PCR and gel electrophoresis to confirm the existence of target sequences.

**Figure 7.**
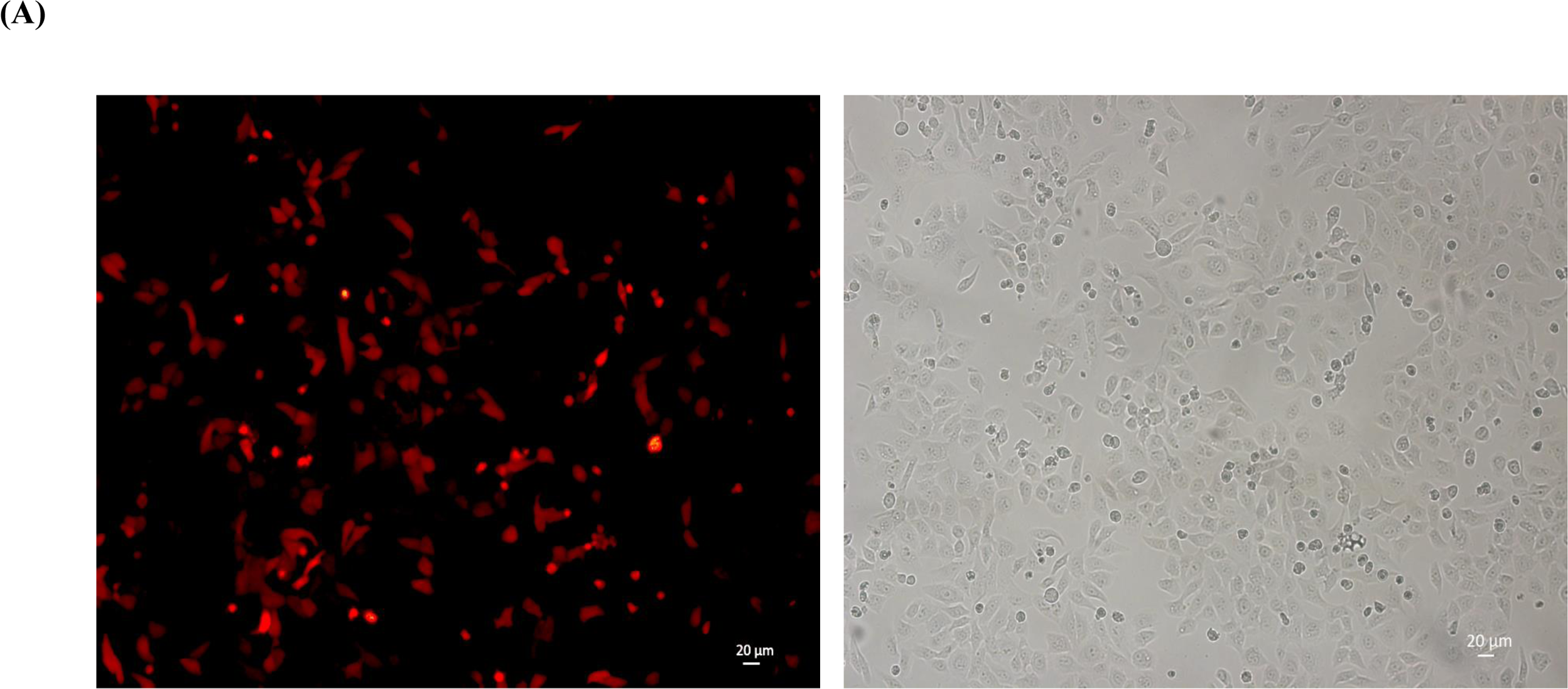
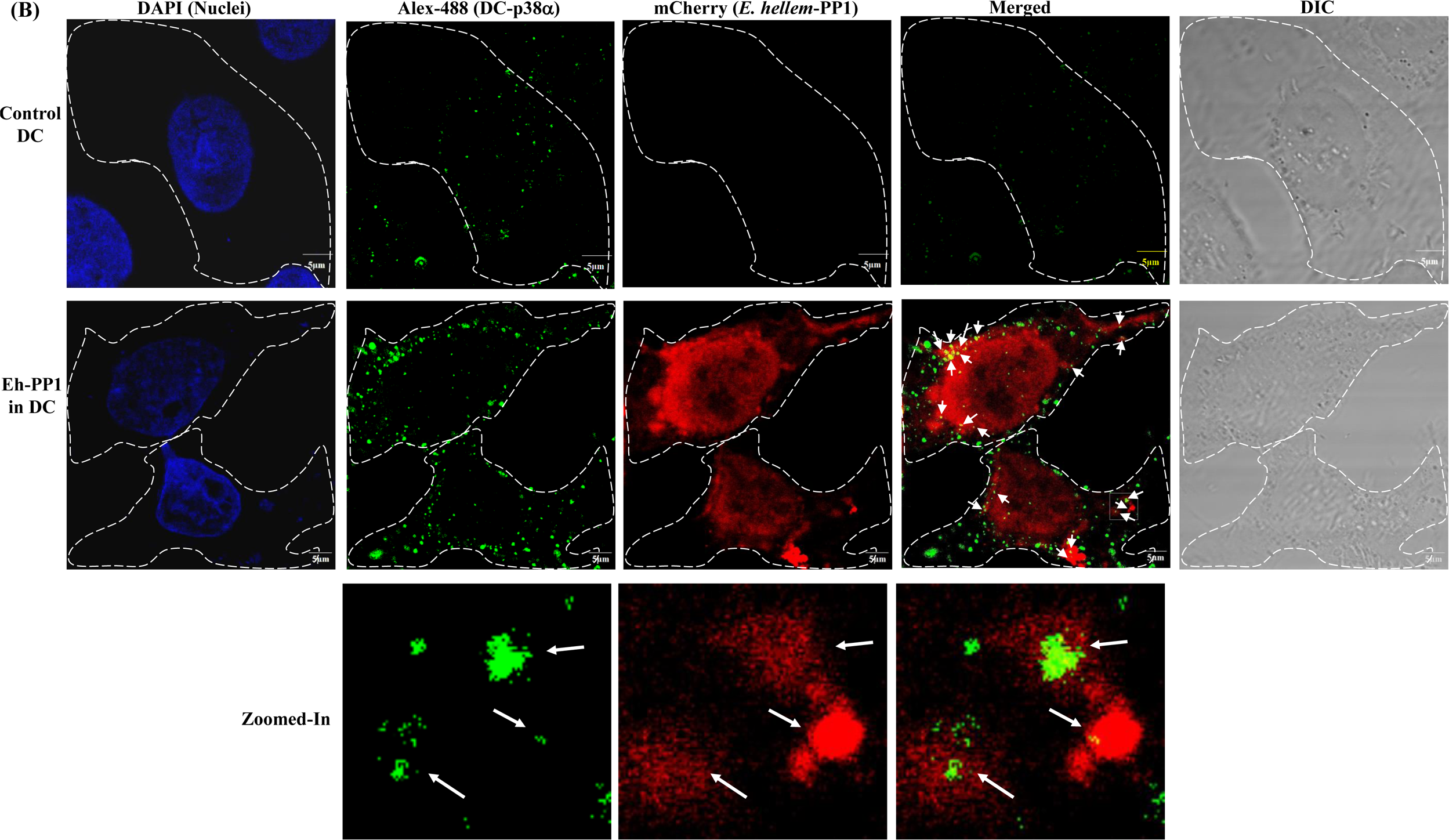
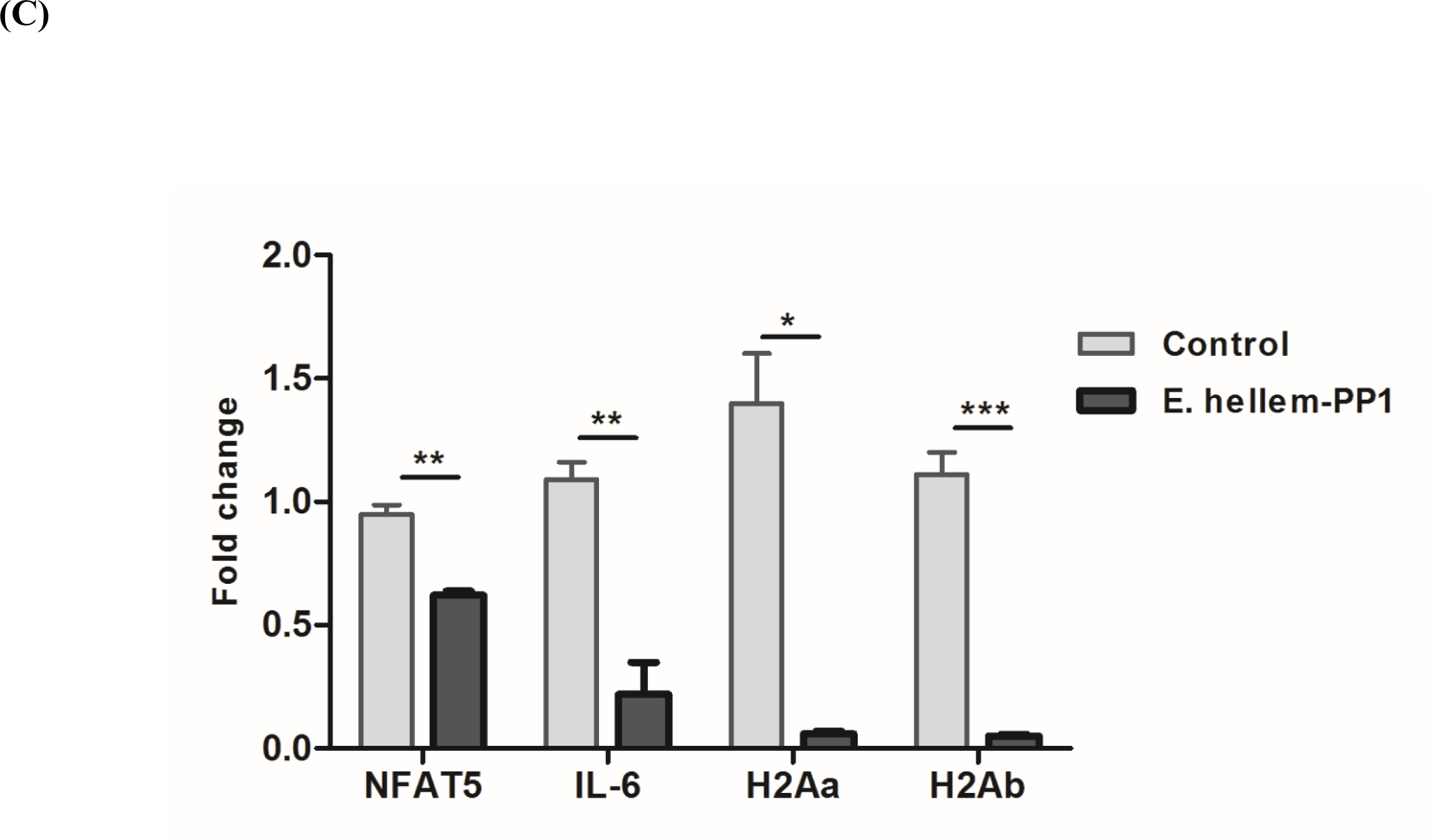
Direct interaction of *E. hellem*-PP1 with DC-p38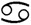 (MAPK14) and the down-regulated MAPK pathway genes expressions. **(A)** Expression of ectopically expressed *E. hellem*-PP1 in DCs. Fluorescent microscopy confirmed the expression of ectopically expressed *E. hellem* derived PP1 (pCMV-mCherry-PP1) in the cytoplasm of DCs (red color). (Scale bar=20 ❍m). **(B)** Immunofluorescent microscopy confirmed the co-localization of *E. hellem*-PP1 with p38α (MAPK14) in DCs. Ectopically expressed *E. hellem* PP1 (mCherry, red color) is expressed in the cytoplasm of DCs. p38α (MAPK14) is labeled with Alex Flour 488 (green color) and is constitutively expressed in the cytoplasm of DCs. Co-localization of *E. hellem*-PP1 with p38α (MAPK14) is indicated with white arrows. Scale bar=5 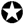m. **(C)** qPCR analysis of *NFAT5, IL-6, H2Aa* and *H2Ab* of MHCII complex genes in either control DCs or DCs expressing ectopic *E. hellem* PP1. All genes tested showed significant down-regulation upon expression of ectopically expressed *E. hellem* PP1. n=12/group; *=P<0.05, **=P<0.01, ***=P<0.001.

Taken together, this is the first clear evidence of a microsporidia-derived protein, E.hellem-derived PP1, directly targeting and modulate host MAPK signaling pathway, thus affecting host immune cell functions. The impaired immune functions of DCs affect both innate and adaptive immune responses and, thus, makes the host more vulnerable for further pathogen infections and co-infections (**Fig. 8**).

**Figure 8.**
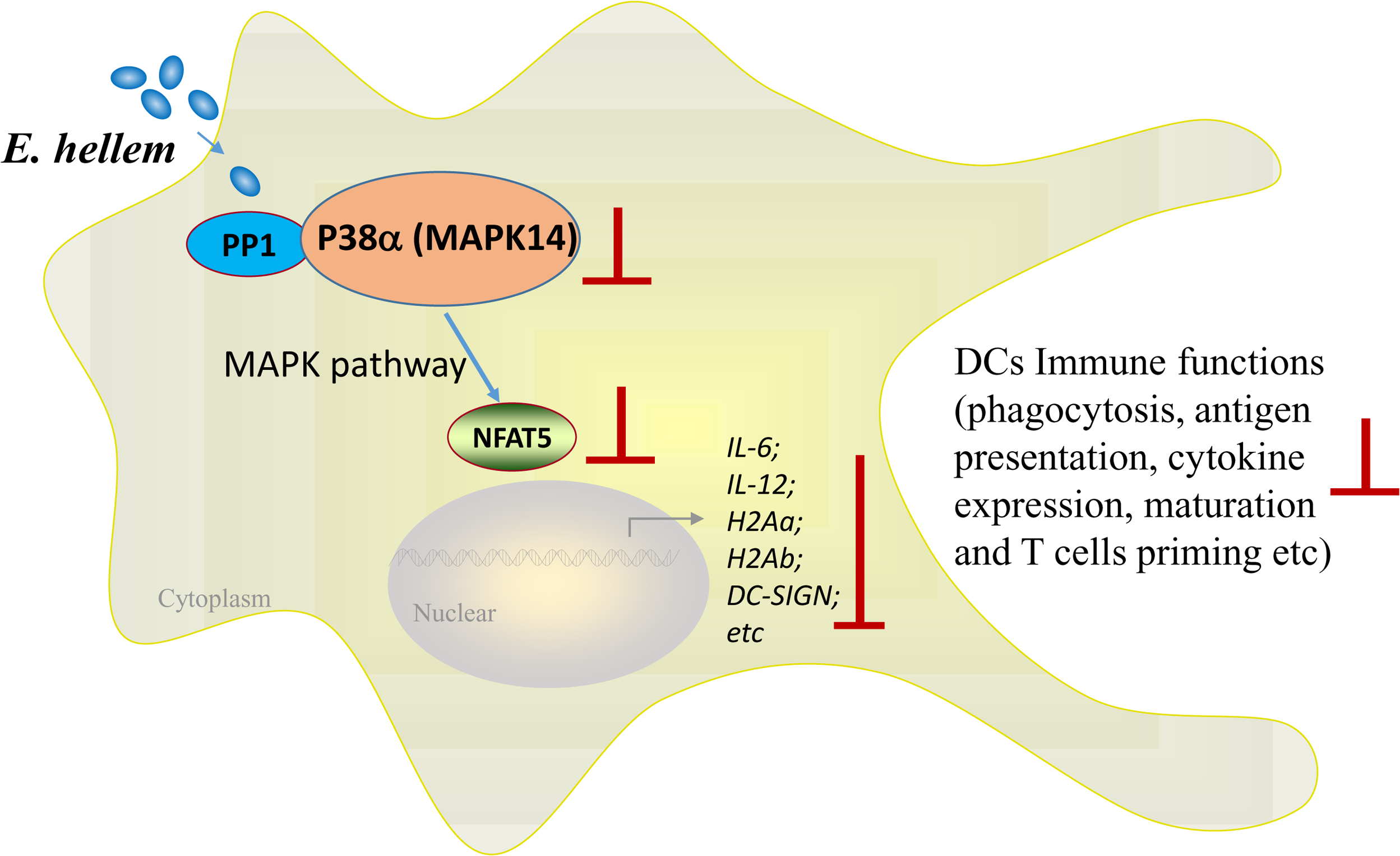
Illustration and summary of proposed mechanism of *E. hellem* disruption of host immune response. *E. hellem* expresses the serine/threonine protein phosphatase PP1 (PP1), which directly interacts with MAPK14 in *E. hellem*-infected DCs. MAPK14 normal functions are then perturbed, leading to downstream downregulation of various genes of the MAPK pathway, such as IL-6, IL-12, DC-SIGN, H2Aa, H2Ab, etc, via disruption of NFAT5. MAPK pathway interference leads to downregulation of other various immune related genes in DCs. The infected DCs are now severely impaired and, since DCs are a lynchpin in the immune response, this leads to disruption of other aspects of both the innate and adaptive immune response in *E. hellem*-infected hosts.

## Discussion

Our study is the first to elucidate the molecular mechanism of microsporidia manipulation of host dendritic cells, and, more importantly, to demonstrate the damaging effects of microsporidia persistence on the immune systems and the subsequent increased chances of being co-infected by other pathogens. In this study, we used the frequently diagnosed and zoonotic species of microsporidia, *E. hellem*, as the pathogen. In the mesenteric lymph nodes of *E. hellem*-infected C57BL/6 mice, we found that dendritic cells (DCs), but not monocytes nor lymphocytes, were the most affected immune cell group. Analysis of immune cells in the spleen further confirmed that the immune function of DCs immune functions were severely impaired after *E. hellem* infection, as reflected by their decreased phagocytic ability, maturation, cytokine expression, and T cell priming potential. *In vitro* and *in vivo* infection models both confirmed that the *E. hellem* infection not only stunted the priming process of infected DCs towards T-cells, but also made infected mice more vulnerable to further infections, as reflected by more weight change and slower recovery rate. To identify the cellular protein(s) and networks perturbed in DCs by *E. hellem* infection, mass spectrometry and yeast two-hybrid analysis were applied; we discovered that the p38α(MAPK14)/NFAT-5 axis of the MAPK signaling pathway in DCs was the key mediator, and *E. hellem* serine/threonine protein phosphatase PP1 directly interacts with p38α(MAPK14) to manipulate DCs immune functions. Knock-down of NFAT5, involved in the p38α(MAPK14)/NFAT-5 axis, via RNAi lead to increased proliferation of *E. hellem* within host, confirming the essential role of MAPK signaling pathways during pathogen-host interactions and providing possible targets of disease prevention and control.

The MAPK signaling pathway is known for its important roles in innate immune responses by regulating the production of inflammatory and anti-inflammatory cytokines, innate immune cell viability, and the function of antigen-presenting cells (APCs) (40). We demonstrated by mass spectrometry that many proteinases in the pathway were disturbed after *E. hellem* infection, such as p38α(MAPK14), MAPK1, Map2k3; many of these perturbed proteins cross-interact with each other. Therefore, *E. hellem* could target any of them to manipulate the whole signal transduction pathway. In fact, we showed the direct interaction between pathogen-derived Serine/threonine-protein phosphatase 1 (PP1) with p38α(MAPK14). This direct interaction indeed impaired the signal transduction process with the MAPK pathway, as we demonstrated in this study that the expressions of transcriptional factors and pro-inflammatory genes/cytokines of DCs were severely detained or inhibited in *E. hellem* infected cells. Besides the proteins in the MAPK pathway, we also identified other differentially expressed proteins such as Neurabin-2, ribosomal protein S6 kinase alpha-1, E3 ubiquitin-protein ligase TRAF7, and others These proteins function as scaffold proteins or act in other cellular pathways such as ERK pathway and ubiquitination (41–43). Thus, it is reasonable to infer that these different signaling pathways were cross-linked together in responding to *E. hellem* infection and modulation. In addition to pathway cross-talk, we are also very interested to identify which pattern recognition receptors were responsible for reception of *E. hellem* and activation of upstream MAPK signaling. Our mass spectrometry analysis provided some candidates for this upstream activation, such as the Integrin-linked protein kinase, which is an adaptor of integrin-related signal reception and transduction (44).

The essential roles and the manipulations of host DC proteins during *E. hellem* infection have been elucidated in our study. Yet, we should not neglect the involvement of other immune cells and processes. For instance, the involvement and protective roles of CD8+ T cells against microsporidia infection have been demonstrated in a mouse model study (45). These findings were in accordance with our findings that, when the T cell priming capabilities were disrupted in DCs, less effector T cells, such as CD8+ T-cells, would be stimulated, therefore, leading to worst outcomes for the host from *E. hellem* infection, along with increased susceptibility to co-infecting secondary pathogens. In addition, the infection of microsporidia in other immune cells may also manipulate normal cell functions. For instance, it is known that p38α(MAPK14) regulates oncogenic processes such as autophagy and apoptosis in many cell types including inflammatory monocytes and B lymphocytes (46, 47). The microsporidia persistence within those cells may not affect immune responses as they do in DCs, but infection in these cells may cause other complications such as increased autophagy or apoptosis, leading to future compromise of the immune system of an infected host. Therefore, it is of great importance to fully investigate the effects of microsporidia on host cells to prevent microsporidiosis and consequent co-infections. Serine/threonine-protein phosphatase 1 (PP1) is a highly conserved protein phosphatase in all eukaryotes, which regulates critical cellular processes including cell cycle progression, apoptosis and metabolism. In mammalian cells, the involvement of PP1 in several oncogenic pathways has become evident, and has been recognized as a potential drug target in cancer (48). In eukaryotic pathogens, the important regulation roles of PP1 are attracting more attention in recent years. For instance, PP1 has been found to regulate pathogen cell maturation, proliferation and metabolism (49, 50). As a result, the existence of pathogen-derived PP1 within host cells may be of great importance for both the intracellular proliferation of the pathogen as well as host cell manipulation. We are very excited to underscore that our study is the first to have identified the existence of *E. hellem*-derived Serine/threonine-protein phosphatase 1 (PP1) and verified its direct interaction with host p38α (MAPK14) in the MAPK signaling pathway. Interestingly, it is reported that PP1 also interacts with and dephosphorylates RPS6KB1, homologous to our identified ribosomal protein S6 kinase alpha-1. Considering the intrinsic cross-talk and links of these *E. hellem*-derived PP1 disrupted host proteins, it will be not surprising to identify multiple effects of microsporidia derived PP1 on host immune responses and other cellular processes in future studies.

In the era of wide-spreading emerging pathogens, shortage of antibiotics and aging of populations, the co-existence and co-infections of multiple pathogens in various populations are becoming great public health threats. Microsporidia exist widely in nature and are asymptomatic during the persist infection in hosts. Our study though was the first to point out the damaging outcomes due to the persistence of microsporidia by impairment of host innate as well as adaptive immunities, thereby increasing host susceptibility to co-infections. Therefore, the detection, prevention, and control of microsporidia should get more attention in the future.

## Author Contributions

JB designed the study and conducted experiments, interpreted the data, and wrote the manuscript. JJ, YT, XW, YC, LC, GA, BM, HZ participated in conducting experiments and analysis of data. GC contributed to study design and manuscript editing. GP and ZY contributed in manuscript editing. All authors contributed to the article and approved the submitted version.

## Acknowledgments

We thank Dr. Louis Weiss, Professor of Albert Einstein College of Medicine; Dr. Bing Han, Professor of Shangdong University, for providing *E. hellem*. Dr. Xiancai Rao, Professor of Department of Microbiology, Army Medical University, for providing EGFP-conjugated *S. aureus*. We also appreciate Dr. Timothy Keiffer, Postdoctoral Fellow at LSU Health Shreveport, for this contributions in correcting the language in this manuscript.

