## Supplementary figures and images for "Microsporidia Ser/Thr Protein Phosphatase PP1 Targets DCs MAPK Pathway and Impairs Immune Functions"

### Supplemental figure1

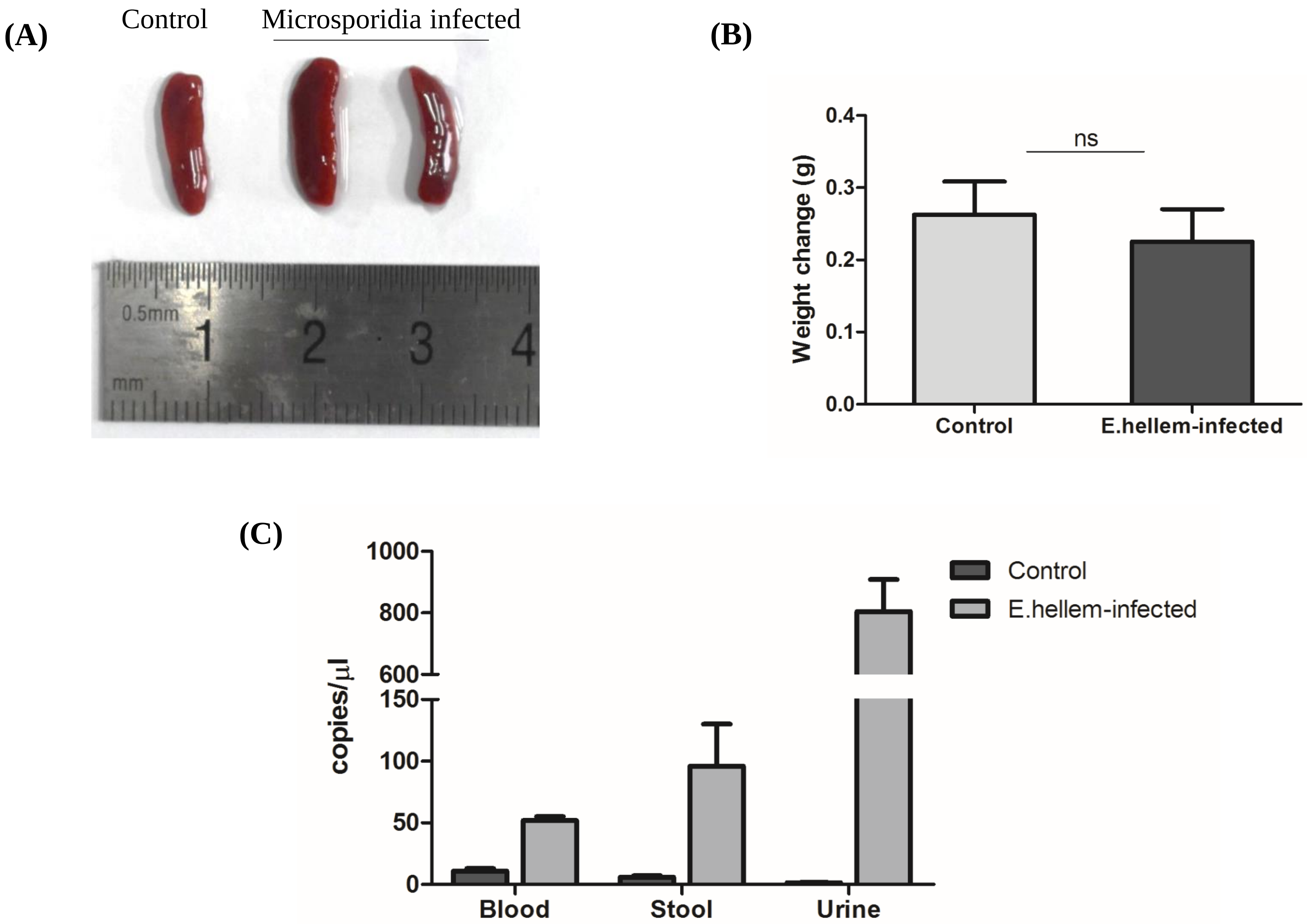
