## supplemental table1 for "Microsporidia Ser/Thr Protein Phosphatase PP1 Targets DCs MAPK Pathway and Impairs Immune Functions"

**S-Table 1.** Primers for qPCR analysis

| **Gene** | **Forward (5′-3′)** | **Reverse (5′-3′)** |
| --- | --- | --- |
| *Nfat5* | TGCTTTCTCAGCTTACCACGG | GTCCGCACAACATAGGGCTC |
| *dc-sign/CD209* | GAGGCTACACTTGGATGGG | AGGGCAGGAAGTTGAAAGC |
| *H2-Aa* | CCTGTGCTGCTGGGTCA | AAGGGATGAAGGTGAGATAAGAC |
| *IL-6* | TAGTCCTTCCTACCCCAATTTCC | TTGGTCCTTAGCCACTCCTTC |
| *IL-12p40* | ATGGAGTCATAGGCTCTGGAAA | CCGGAGTAATTTGGTGCTTCA |
| *IFNα2* | TGCCTCACACTTATAACCTCAG | GTATTCCAAGCAGCAGATGAAG |
| *IFNγ* | ACAGCAAGGCGAAAAAGGATG | GGTGGACCACTCGGATGA |
| *CD4* | TGCCTCAGTATGCTGGCTCT | GAGACCTTTGCCTCCTTGTTC |
| *PD-1* | CCAGGATGGTTCTTAGACTCCC | TTTAGCACGAAGCTCTCCGAT |
| *TIM-3* | GAGTTACGGGACTCTAGATTGG | TGTTTTCTTCTGAGCGAATTCC |
| *CTLA* | TTCTCTTCATCCCTGTCTTCTG | CTCATTCCCCATCATGTAGGTT |
| *Human β-actin* | CATGTACGTTGCTATCCAGGC | CTCCTTAATGTCACGCACGAT |
| *Mouse β-actin* | GCTGTCCCTGTATGCCTCTG | TGATGTCACGCACGATTTCC |
